# A focal traumatic injury to the spinal cord causes an immediate and massive spreading depolarization sustained by chloride ions, with transient network dysfunction and remote cortical glia changes

**DOI:** 10.1101/2024.07.15.603535

**Authors:** Atiyeh Mohammadshirazi, Graciela L. Mazzone, Benjamín A. Zylberberg, Luca Mio, Giulio Pistorio, Carmen Falcone, Giuliano Taccola

## Abstract

In clinics, physical injuries to the spinal cord cause a temporary motor areflexia below lesion, known as spinal shock. This topic is still underexplored due to the lack of preclinical SCI models that do not use anesthesia, which would affect spinal excitability. Our innovative design considered a custom-made micro impactor that provides localized and calibrated strikes to the ventral surface of the thoracic spinal cord of the entire CNS isolated from neonatal rats. Before and after injury, multiple ventral root (VR) recordings continuously traced respiratory rhythm, baseline spontaneous activities, and electrically-induced reflex responses. As early as 200 ms after impact, an immediate transient depolarization spread from the injury site to the whole spinal cord with distinct segmental velocities. Stronger strikes induced higher potentials causing, at the site of injury, a transient drop in tissue oxygen levels and a massive cell death with complete disconnection of longitudinal tracts. Below the impact site, expiratory rhythm and spontaneous lumbar activity were suppressed. On lumbar VRs, reflex responses transiently halted but later recovered to control values, while electrically-induced fictive locomotion remained perturbed. Moreover, low-ion modified Krebs solutions differently influenced impact-induced depolarizations, the magnitude of which amplified in low-Cl^−^. Moreover, remote changes in cortical glia occurred soon after spinal damage. Overall, our novel in vitro platform traces the immediate functional consequences of impacts to the spinal cord during development. This basic study provides insights on the SCI pathophysiology, unveiling an immediate chloride dysregulation and transient remote glial changes in the cortex.

## Introduction

A spinal cord injury (SCI) demonstrates that the mature central nervous system (CNS) cannot regenerate nor repair itself after traumatic insults. Because of this vulnerability, an SCI often causes a permanent loss of sensory and motor control over the body parts innervated by spinal neurons located below the level of injury. Eventually, an SCI results in a long-life debilitating condition characterized by motor paralysis and a variegated spectrum of functional deficits and complications. To date, there is no cure against paralysis and current rehabilitation still focuses mainly on strengthening the able part of the body to compensate for the loss of volitional motor control over the rest. Support in daily tasks mainly occur through classical mobility aids, such as a wheelchair and crutches, but also using newly introduced technologies, such as exoskeletons (Gad et al., 2017) and advanced brain machine interfaces (Lorach et al., 2023), which however allow only minor functional benefits.

Nevertheless, some scattered and unpredictable spontaneous neurologic recoveries have been reported (Kirshblum et al., 2021) and, in less severe injuries, a substantial spontaneous regain of functions plateaued at 16 weeks after injury (Geisler et al., 2001). Spontaneous recoveries still challenge our understanding of the pathophysiological mechanisms of an SCI and of the residual potential of the cord to repair spinal circuits.

In particular, pediatric spinal injuries, which account for 1-10% of all SCIs (Carreon et al., 2004), show higher rates of spontaneous functional recovery compared to adults (Eleraky et al., 2000; Wang et al., 2004). Likewise, the study of traumatic injuries in the developing mammalian spinal tissue, also in comparison with adults (Clarke et al., 2009), is compelling to clarify the peculiar pathophysiological mechanisms of neonatal SCIs, in the hope to identify the reasons for the enhanced recoveries in children and possibly expand them to all people with SCI.

Established models of SCI use adult mammals (Kjell and Olson, 2016) under anesthesia. However, when administered near the time of injury, anesthetics affect the damage progression as, based upon the different drug adopted, they can exert a neuroprotective effect (Salzman et al., 1993; Davis and Grau, 2023) or, on the contrary, exacerbate the hypoxic neuronal injury caused by transitory hypotension (Robba et al., 2017). In addition, up to date, only few reports described standardized and calibrated SCI models using immature spinal tissues (Taccola et al., 2010; Mladinic et al., 2013).

Another missing tile for the overall understanding of an SCI is the identification of the immediate events that take place during a physical impact to the spinal cord. In particular, it is still unknown how the primary mechanical insult to the spinal tissue contributes to trigger the subsequent cascade of pathological events known as secondary damage, which eventually determines the extent of tissue damage and hinders the chances of achieving a functional recovery (Carlson et al., 1998).

Indeed, after injury, a temporary loss or depression of all, or most, spinal reflex activity takes place below the lesion. This phenomenon is called spinal shock, and the underlying mechanisms are not fully clarified (Ditunno et al., 2004). A spinal shock clinically persists for days or weeks, depending on which reflex is clinically being tested for reappearance. However, when duration is defined based on the initial recovery of any one reflex, then the spinal shock lasts no longer than 20 - 60 min (Ditunno et al., 2004).

The lack of reflex activity has been mainly attributed both to the sudden disappearance of the predominantly facilitatory tonic influence exerted by descending supraspinal tracts, and to an increased presynaptic inhibition. In addition, depression of synaptic activities also depends on the hyperpolarization of spinal neurons due to an excessive accumulation of potassium (Atkinson and Atkinson, 1996).

In pediatric SCIs, reflexes recover sooner, likely because descending supraspinal tracts in children are not fully developed, thereby normal descending inhibition to spinal inhibitory pathways is less affected by an SCI compared to adults, mitigating the depression of spinal networks during shock (Guttmann, 1976).

Experimentally, the main features of a spinal shock parallel those of an early depolarization of the entire spinal cord following a trauma, also known as injury potential, which spreads rostrally and caudally from the site of impact. This early depolarization is sustained by a transient extracellular ionic disbalance (Goodman et al., 1985; Wang et al., 2015) and is similar to a cortical spreading depression (SD), which exhibits a marked, enduring reduction in the intrinsic electrical activity of neurons, eventually spreading from the original source out in all directions and involving increasingly distant parts of the cerebral cortex (Leao, 1944, 1947). A cortical SD is triggered, among other causes (Gerasimova et al., 2021), by traumatic brain injuries (Hermann et al., 1999).

In both amphibians and rodents, a compressive injury to the cord is followed both by SD-like waves characterized by a velocity of propagation of around 10-15 mm/min, and by a rapid and reversible increase in extracellular concentrations of K^+^ ions (Streit et al., 1995; Gorji et al., 2004). Interestingly, electrically evoked potentials were transiently abolished during spinal SD waves, eventually returning to baseline values only after about twenty minutes. This phenomenon suggests that spinal SD might determine areflexia after spinal shock (Gorji et al., 2004). The same study also described how the SD evoked by an injury to the brain cortex reduced excitability of spinal neurons located in upper spinal segments, indicating that SD-like waves induced by an injury maintain a form of conduction among cortical and spinal structures (Gorji et al., 2004).

However, the appearance of any neuronal changes in the brain after SCI is controversial, with conclusions spanning from the absence of cellular loss (Crawley et al., 2004) to extensive retrograde neurodegeneration (Feringa and Vahlsing, 1985; Hains et al., 2003). A detailed study after spinal cord contusion in mice described SCIs as complex events affecting the entire CNS and generating cognitive changes and depressive-like behaviors, associated with reactive microglia and neuronal loss in the hippocampus and cerebral cortex (Wu et al., 2014). However, the authors failed to detect a significant neuronal death in brain districts even after two weeks after SCI, suggesting only chronic inflammatory changes. However, it is still unknown whether immediate pathological signals are transiently triggered in the brain right after an impact to the spinal cord. The presence of a pathological sign could actually provide a novel marker to more realistically characterize the severity of a lesion and envisage potential recoveries. Obstacles to the comprehension of a spinal shock and the related transient changes in the brain reside in some technical challenges that arise from the preclinical models currently available. Indeed, fully anesthetized animals do not allow to record the electrical activity of spinal neurons in the same instants when the physical impact occurs, due to both motion and electrical artifacts generated by standard experimental impactors, which interfere with the low amplitude of currents involved. As a consequence, the earliest injury potential has been recorded only after four minutes from the impact (Goodman et al., 1985). This temporal limitation sums up to the effects of anesthetics that depress neuronal excitability and are used in preclinical models at the time of the physical trauma. A solution to avoid any technical artifacts, as well as any consequences of anesthetics, is the adoption of the neonatal preparation of the entire central nervous system in vitro (CNS; (Mohammadshirazi et al., 2023; Apicella and Taccola, 2023)), which does not require the administration of any drugs. In addition, the rodent spinal cord can be optimally damaged through a low-noise calibrated micro impactor recently designed in the laboratory.

Using this setting, we aim at quantifying the immediate events triggered by a physical insult to the cord and their spread from the site of injury both caudally toward spinal segments, and rostrally up to brain structures. In addition, we will assess any spontaneous functional recoveries occurring in the neonatal spinal circuitry.

## Methods

### In vitro preparation of the isolated entire CNS

All procedures were approved by the International School for Advanced Studies (SISSA) ethics committee and are in accordance with the guidelines of the National Institutes of Health (NIH) and the Italian Animal Welfare Act 24/3/2014 n. 26, implementing the European Union directive on animal experimentation (2010/63/EU). Every effort was made to minimize the number of animals used and to ensure their well-being. A total of 92 postnatal (P) Wistar rats (P0-P3) of random sexes were included in this study.

Experiments were performed on in vitro preparations of the entire isolated central nervous system (CNS; (Mohammadshirazi et al., 2023)). Newborn rats were subjected to cryoanesthesia (Danneman and Mandrell, 1997). After disappearance of the paw pinch reflex, surgical procedures considered the quick removal of: forehead at orbital line, ribs cage, internal stomach and forelimbs. The preparation was then transferred to a petri dish filled with oxygenated Krebs solution that contained (in mM): 113 NaCl, 4.5 KCl, 1 MgCl27H2O, 2 CaCl2, 1 NaH2PO4, 25 NaHCO3, and 30 glucose, gassed with 95% O2 - 5% CO2 (PO2 533.65 ± 44.05 Torr), pH 7.4, 299.62 ± 3.2 mOsm/kg. Under microscopic guidance, craniotomy and ventral laminectomy were performed keeping the dorsal vertebra and dorsal root ganglia (DRG) intact. Afterwards, the entire CNS preparation was maintained in oxygenated Krebs solution at room temperature for 15 minutes and then mounted in the recording chamber (total volume = 4.7 ml, flow rate = 7 mL/min, controlled temperature = 25-27ᵒ C, TC-324C Warner Instruments, USA). For stable electrophysiological recordings, the preparation was fixed ventral side up with insect pins passing through dorsal vertebrae. For selective root recordings, VRs and DRs were detached from DRGs.

### Extracellular Recordings

DC-coupled recordings were obtained from both VRs and DRs using monopolar suction electrodes realized by pulling tight-fitting glass pipettes (1.5 mm outer diameter, 0.225 mm wall thickness; Hilgenberg, Germany). Electrodes were connected to a differential amplifier (DP-304, Warner Instruments, Hamden, CT, USA; high-pass filter = 0.1 Hz, low-pass filter = 10 kHz, gain X 1000). Analog signals were filtered through a noise eliminator (D400, Digitimer Ltd, UK), then digitized (Digidata 1440, Molecular Devices Corporation, Downingtown, PA, USA; digital Bessel low-pass filter at 10 Hz; sampling rate = 5 kHz) and visualized real-time with the software Clampex 10.7 (Molecular Devices Corporation, Downingtown, PA, USA).

### Electrical Stimulation

Trains of rectangular electrical pulses (pulse duration = 0.1 ms, frequency = 0.1 Hz) were supplied to sacrocaudal afferents through a programmable stimulator (STG4002, Multichannel System, Reutlingen, Germany) using bipolar glass suction electrodes connected two close silver wires (500-300 µm). Stimulus intensity (40-160 µA) was attributed as times to threshold (Th), where Th is the lowest intensity required to elicit a small deflection of VRrL5 baseline. To generate epochs of fictive locomotor patterns (Kiehn, 2006), 160 rectangular pulses (duration = 0.1 ms, intensity = 37.5-150 µA, 1-5 × Th) were supplied at 2 Hz to sacrocaudal afferents for a total length of 80 s. Recordings were acquired in the same preparation from r and l VRL2 (for bilateral flexor commands) and VRrL5 (for extensor output).

### Spinal cord injury

A calibrated physical impact to the thoracic (T) cord was provided using a custom-made micro-impactor device specifically designed and shielded to allow simultaneous electrophysiological recordings from the neonatal CNS in vitro. The device is currently being patented by SISSA and is available upon request (https://www.valorisation.sissa.it/device-mechanically-stimulating-biological-material-and-its-procedure). The impactor tip (diameter = 2 mm) was precisely positioned on the ventral surface of the spinal cord (T10) using a manipulator. The micro-impactor was controlled through a dedicated software that allows to precisely set impact parameters (displacement, speed, acceleration, deceleration, and pause time). In our experiments, the maximum severity of compression without completely transecting the neonatal spinal cord (diameter around 3-4 mm) was obtained with the impactor tip descending into the cord by 2656 µm from the spinal surface, at an average speed of 4 mm/s, maintaining an acceleration and deceleration of 6.1 ± 0.05 mm/s^2^. After the impact, the tip of the impactor was promptly returned to its original position at the same speed, acceleration, and deceleration. For minor severities of injury, varying displacements and velocities of the impactor rod were considered (625 µm at 2 mm/s, 1250 µm at 2.8 mm/s, and 1875 µm at 3.4 mm/s), while keeping acceleration and deceleration constant.

### Modified Krebs solutions

Three modified Krebs solutions were prepared. The low chloride solution (in mM) was composed of: 56.5 NaCl, 56.5 sodium isethionate, 4.5 KCl, 1 MgCl27H2O, 2 CaCl2, 1 NaH2PO4, 25 NaHCO3, and 30 glucose (297.62 ± 3.8 mOsm/kg). The low calcium solution (in mM) contained: 113 NaCl, 4.5 KCl, 1 MgCl27H2O, 1 CaCl2, 1 NaH2PO4, 25 NaHCO3, and 30 glucose. The low potassium solution (in mM) was prepared with: 113 NaCl, 2.25 KCl, 1 MgCl27H2O, 2 CaCl2, 1 NaH2PO4, 25 NaHCO3, and 30 glucose. The three modified Krebs solutions were gassed with 95% O2 - 5% CO2 and their osmolarity was adjusted by adding sucrose to mimic the osmolarity of control Krebs solution. To assess the different impact of modified Krebs solutions on spinal reflexes, analysis was performed on five responses randomly chosen in control, and five in the last two minutes of low-ion perfusions (from the 88^th^ to 90^th^ minute for low Cl^−^, and from the 28^th^ to 30^th^ minute for low Ca^2+^ and low K^+^).

### Tissue Oxygen Assessment

PO2 measurements in the spinal cord were conducted using a fiber-optic microsensor with a 50 μm tip diameter (Optode, OxyMicro System, World Precision Instruments, FL, USA). The microsensor was implanted at 100µm deep into the cord in the anterior funiculus between L1 and L2 segments. Measurements were taken at the sampling rate of one per second and were directly acquired using OxyMicro v7.0.0 software (OxyMicro System, World Precision Instruments, FL, USA). Temperature during all PO2 measurements was maintained within the range of 25-27 °C.

To ascertain whether high K^+^ perfusions affect measurements of the microsensor probe, test experiments considered positioning the tip of the microsensor in the recording chamber void of any preparation. PO2 values remained unchanged in standard Krebs solution (610.29 ± 7.63) and during 10 mM K^+^ applications (613 ± 8.53) indicating that tissue oxygen assessments did not change during perfusion with potassium ions (Sl. Fig. 4 B).

### Slice Immunostaining and Cell Counting

After electrophysiological experiments, spinal cords were fixed overnight in 4% paraformaldehyde at 4°C and then tissue was cryopreserved with 30% sucrose in phosphate buffered saline (PBS), following our standard procedures (Taccola et al., 2008, 2010). Spinal cords were cut in 20-μm (coronal or longitudinal) slices using a sliding cryostat microtome. To detect neurons and motoneurons, slices were incubated overnight at 4 °C with blocking solution for 1 h and then with mouse monoclonal anti-NeuN or SMI 32 antibody (1:200; Merck Millipore, Milan, Italy; ABN78 and 1: 200; Covance, Berkeley, CA, catalog # SMI-32P, respectively) in 5% fetal calf serum (FCS), 5% bovine serum albumin (BSA), and 0.3% Triton X-100 in PBS. After three washes in PBS, floating sections were incubated for 2 h at room temperature with the goat anti-mouse Alexa 488-labeled secondary antibody (1:500, Invitrogen). To visualize cell nuclei, slices were incubated in 1 μg/ml solution of 4’, 6-diamidino-2-phenylindole (DAPI). Sections were washed three times in PBS for 5 min and mounted using Vectashield® medium (Vector Laboratories, Burlingame, CA) and coverslips. NeuN and SMI 32 positive cells were assessed in a complete set of z-stack images, typically at a depth of 4-μm, using confocal series acquired by Nis-Eclipse microscope (20x magnification, NIKON, Amsterdam, Netherlands) at 20× magnification. The number of SMI 32 positive cells was determined by VolocityTM software (https://www.volocity4d.com, Improvision, Coventry, UK) while Image J software (NIH, https://imagej.nih.gov/ij/index.html) was adopted for NeuN positive cells.

### Cortical glia immunohistochemistry and image acquisition

Brain specimens from 16 pups which fixed at 25 minutes, 1.5 hours, 2.5 hours post-injury were sectioned into 20 µm-thick coronal sections using a cryostat. Sections were processed for free-floating immunohistochemistry following a previously published protocol (Ciani et al., 2023). In detail, slices were pre-treated with 0.1% Sudan Black (in 70% Ethanol) for 30 min for autofluorescence reduction, quickly washed with 70% ethanol, and incubated with 10% serum blocking solution for 1 h at room temperature. Then, sections were incubated with the primary antibody, anti-S100b (rabbit, Abcam #ab52642, RRID: AB_882426), diluted in 2% blocking solution and stored overnight at 4° C degrees. Subsequently, sections were washed with 1X-PBS three times, and incubated with the secondary antibody, AlexaFluor#488 conjugated polyclonal anti-rabbit antibody (1:400, Invitrogen, #A-21206 RRID: AB_141633), diluted in 2% serum blocking solution. Finally, DAPI was used (1:500, #10236276001, Hoffmann-La Roche, Basel, Switzerland) to stain nuclei.

Pictures were taken using Nikon A1/R confocal, with a 60X oil objective. To include all cells among the whole tissue thickness, images were taken as z-stacks of at least 20 steps of 1 μm, to include an optical section thickness of at least 20 μm.

### Data Analysis

Data analysis was performed using Clampfit 10.7 software (Molecular Devices Corporation, PA, USA). Spontaneous rhythmic motor discharges recorded from cervical VRs, with a period of 24.27 ± 16.13 s, were attributed to respiratory bursts that were also derived synchronous among bilateral lumbar VRs (Mohammadshirazi et al., 2023; Apicella and Taccola, 2023). The coefficient of variation (CV), an indicator of response consistency, was determined by the ratio between standard deviation and mean value (Taccola et al., 2020). To calculate conduction velocity, the latency of each response was divided by the distance between the center of the impacted area and the recording sites, as precisely measured using a microcalibrated dial caliper (sensitivity = 20 µm). The correlation coefficient function (CCF) was used to measure phase coupling between pairs of VRs using Clampfit 10.7 software. A CCF value of ≥ 0.5 indicates synchronous rhythmic signals from two VRs, while a CCF value of ≤ −0.5 indicates alternating signals (Taccola and Nistri, 2005; Dose et al., 2016).

Immunofluorescent images from cerebral cortex were analyzed to quantify the number of both S100b+ astrocytes and total DAPI+ cells, per each region of interest (ROI) using the Volocity software (Quorum Technologies Inc., CA).

### Statistical Analysis

Statistical analysis was performed with GraphPad InStat 3.10 (Inc., San Diego, California, USA). In the Results section, the number of animals is denoted as “n”, and data is presented as mean ± standard deviation (SD) values. Before conducting comparisons among groups, a normality test was performed to select the appropriate parametric or non-parametric tests. Parametric data were analyzed with paired student t-test, one-way analysis of variance (ANOVA) or repeated measure analysis, while non-parametric data were analyzed using Kruskal-Wallis, Mann-Whitney, Friedman, or Wilcoxon matched-pairs signed-ranks tests. Multiple comparisons ANOVA was followed by Tukey-Kramer or Dunnett multiple comparisons tests. Differences were considered statistically significant when P value ≤ 0.05.

## Results

### A physical impact to the cord elicits an immediate depolarizing potential

To investigate the immediate events following a contusive spinal cord injury, a custom-made impactor was employed to induce a physical impact at thoracic spinal cord level of an in vitro preparation of entire CNS (Mohammadshirazi et al., 2023). The careful design of the impactor included a proper shielding to minimize any electrical interference during operation, to allow simultaneous electrophysiological recordings during the impact.

In an exemplar experiment, a brief and intense impact (time = 650 ms, displacement = 2656 µm) on the ventral cord (T10) led to a massive depolarization, simultaneously recorded rostral and caudal to the compression site from cervical and lumbar VRs, respectively (Fig. 1 A). Injury-induced potentials started 194.4 ms after the impact on VRrL5, and 225.2 ms on VRrC2. On VRrL5, a depolarization peak of 6.86 mV is reached after 2.66 s, followed by a depolarizing plateau lasting 3.52 s (Fig. 1 B) and spontaneously recovering to baseline in less than 15 minutes. VRrC2 generated a smaller depolarization peak (1.47 mV). The profile of the average injury-induced potential from VRrL5 reveals a peak of 8.21 ± 1.32 mV and a latency of 178.41 ± 15.17 ms after the impact, recovering to 81.11 ± 12.56 % six min later (Fig. 1 G).

**Figure 1.**
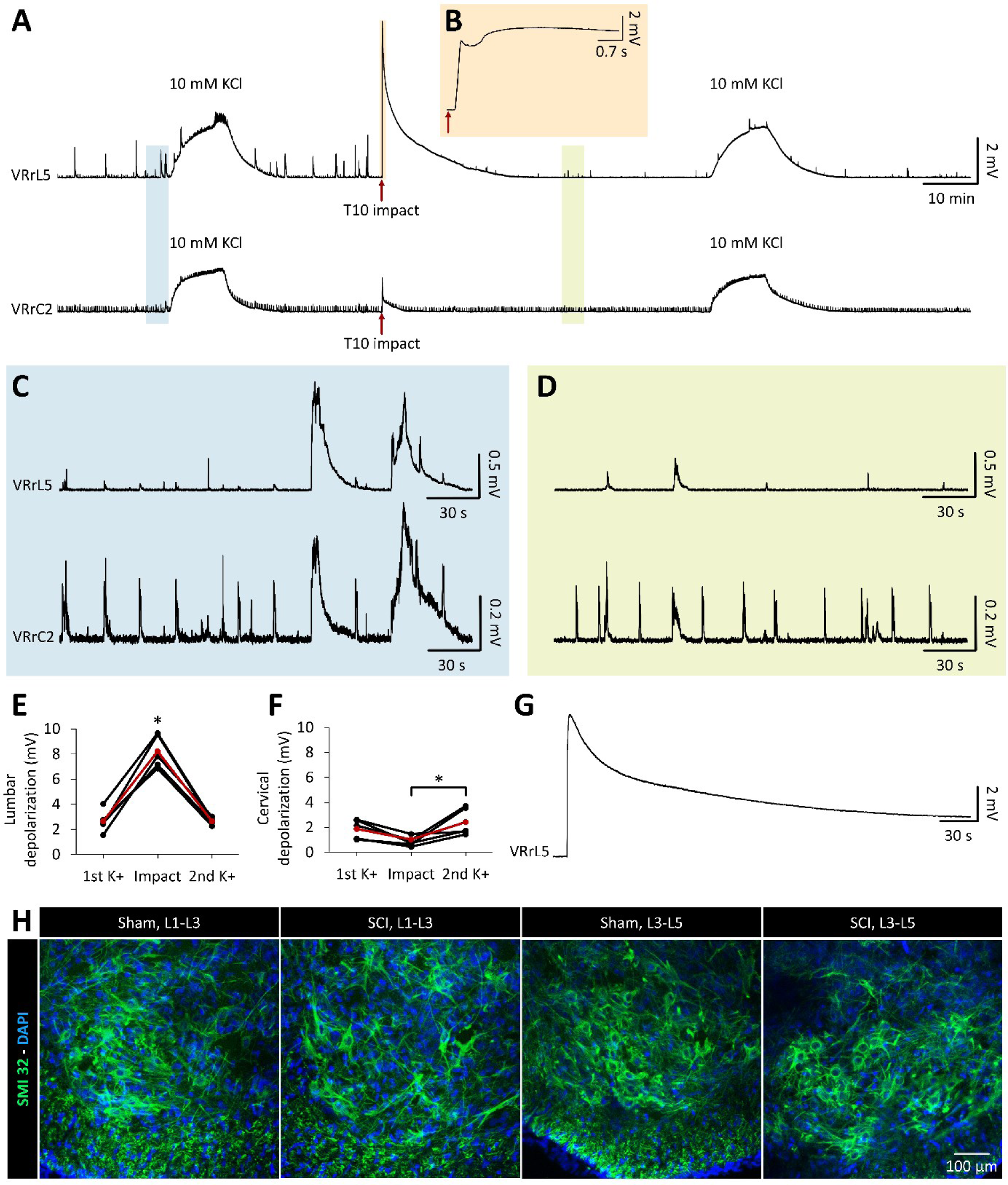
A transient depolarization immediately follows a physical injury to the spinal cord. **A.** Long continuous recordings from VRrL5 and VRrC2, while the cord is being impacted at T10 (red arrows). Before and after the impact, 10mM KCl were perfused for ten minutes to compare maximum recruitment of motor pools. **B.** Magnification highlights the depolarization at VRrL5 in the first five seconds after impact (red arrows). **C, D.** Faster time scales of VR traces in A, corresponding to the shaded blue and green fields that are recorded before and after the impact, respectively. After the impact, spontaneous rhythmic respiratory events on VRrC2 persist unchanged, while the lumbar respiratory activity disappears. Spontaneous sporadic bursting from VRs is largely reduced by the trauma. **E.** From pooled data from five experiments, amplitude of impact-induced depolarization recorded from VRrL5 significantly exceeds the depolarization-induced by 10 mM KCl before and after the impact (*, P < 0.001). Mean values are indicated by the red dots and line. **F.** In cervical motor pools, depolarization after injury is notably smaller than after a second application of potassium (*, P = 0.046). Mean values are indicated by the red dots and line. **G.** Superimposed depolarizations from VRrL5 in five experiments. **H**. SMI 32 labelling of samples collected 90 min after the impact display a comparable number of motoneurons in L1 to L3, and in L3 to L5 segments of both, sham and SCI experiments.

To confirm that the observed sudden increment in DC levels is indeed a genuine potential rather than an artifact, we performed supplementary four experiments, Firstly, where the device solely acted in the bath close to the preparation, without touching the cord. (Sl. Fig. 1 A, B). Furthermore, when multiple impacts of equal severity (displacement = 2656 µm) were serially applied to the same site (T10) for five times, with a lag of less than 10 seconds between any two consecutive impacts, peaks of injury-induced potentials remained stable, hence excluding any summation of artifacts (Sl. Fig. 1 C). In another trial, the impact was delivered at the top of a large depolarization (16.46 mV) produced by perfusing 50 mM KCl. No injury-evoked depolarization was noticed when the preparation was already maximally depolarized by the high K^+^ concentrations (Sl. Fig. 1 D). Finally, no baseline deflections were recorded from VRrL5 when the impact was inflicted to the T10 segment of a spinal tissue inactivated by both high temperature (100ᵒ C) and long-lasting (1 h) oxygen deprivation (Sl. Fig. 4 C), proving the biological origin of depolarization after injury. Collectively, these tests revealed the absence of any baseline drift produced either by the engine itself or by the sudden movement of the tip in the recording bath.

To monitor the respiratory rhythm originated by neuronal networks located in the brainstem (Del Negro et al., 2018), spontaneous rhythmic bursts were recorded from cervical VRs of the isolated CNS (Mohammadshirazi et al., 2023; Apicella and Taccola, 2023). The respiratory rhythm can also be recorded from lumbar VRs, which drive the recruitment of chest muscles to assist the expiratory phase (Giraudin et al., 2008). Noteworthy, respiratory bursting recorded from upper cervical VRs, 30 mins after injury, was not affected by the thoracic impact to the cord (Fig. 1 C, D). In seven preparations, respiration frequency from VRC2 was 84.28 ± 20.29 % of pre-impact control (P = 0.709, paired t-test). To assess any early and transient alteration of the respiratory rhythm during the impact, 20 respiratory bursts from cervical VRs were analyzed right before and soon after the injury. In 4 out of 7 preparations, the first respiratory event after the impact was delayed, showing an early perturbation of the neuronal networks in the brainstem generating the respiratory rhythm (Sl. Fig. 2). Albeit not consistent among all preparations, this effect was observed in the majority of experiments, regardless of the magnitude of injury potentials from cervical VRs and the age of animals (Sl. Fig. 2).

**Figure 2.**
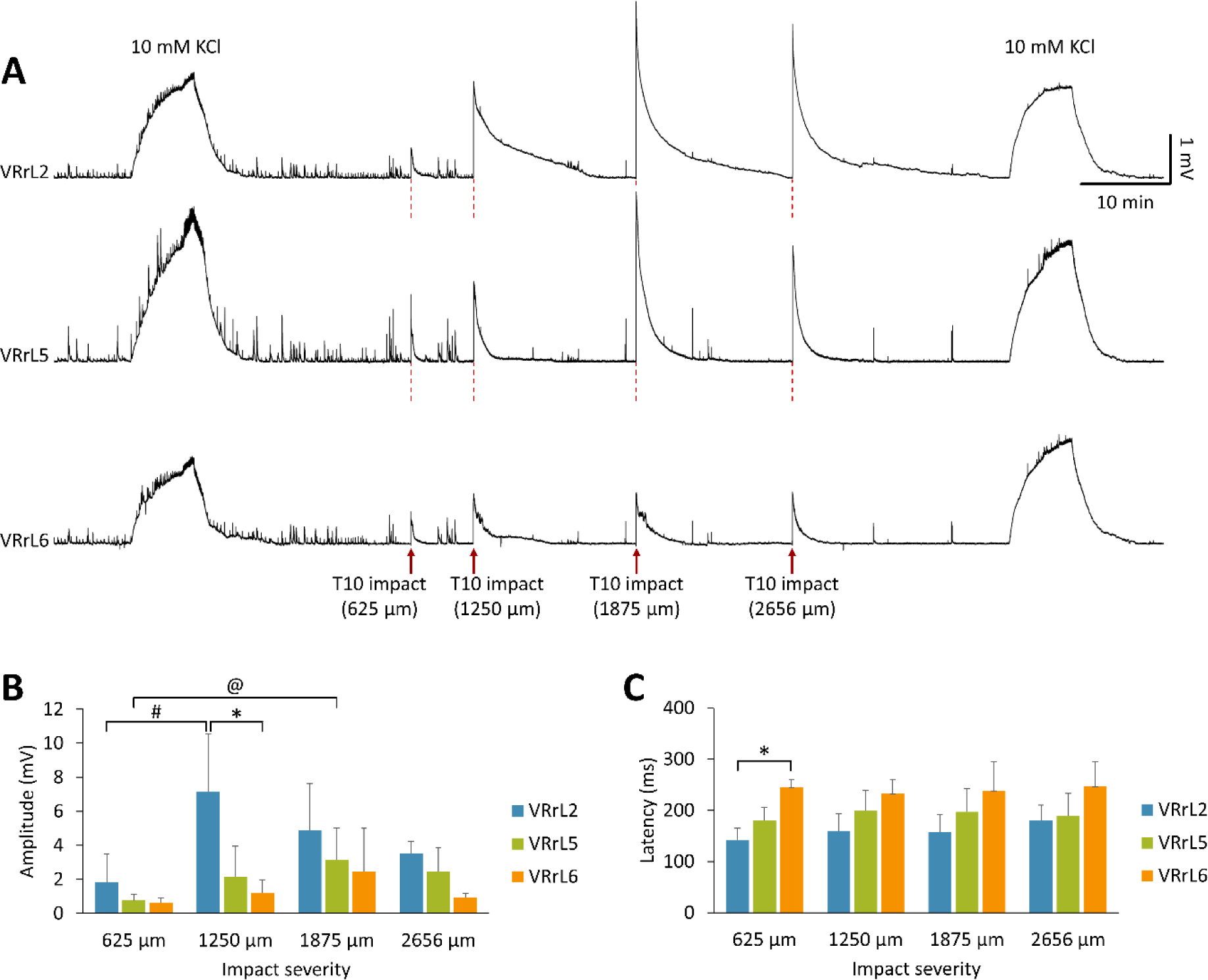
Calibrated impacts of increasing severity elicit higher injury potentials and dramatically reduce spontaneous network activity. **A.** Simultaneous recordings from VRrL2, VRrL5, VRrL6, during four serial impacts of increasing severity (from 625 to 2656 µm of tip displacement) at T10. Spontaneous baseline activity progressively reduces with stronger impacts, until its almost complete suppression after repetitive injuries. At the beginning and at the end of the experiment, 10 mM potassium were administered to confirm the unaffected recruitment of motor pools. **B.** Bars summarize the amplitude of injury-induced depolarizations arranged by different severities of impact. Larger injury potentials appear from VRs closer to the injury site (*P = 0.009 for VRrL2 vs. VRrL6 at 1250 µm) and after impacts of increasing strengths (^@^P = 0.019 for 625 µm vs. 1875 µm from VRrL5; ^#^P = 0.024 for 625 µm vs. 1250 µm from VRrL2). **C.** Histogram reports the latency of injury-induced depolarizations for different severities of impact. The injury potential spreads from the lesion site to lumbar segments following a rostrocaudal propagation (*P = 0.001, for VRrL2 vs. VRrL6).

Contrariwise, impact at T10 largely suppressed on both VRrL5 and VRrC2 those sporadic episodes that appeared synchronous among all neonatal motor pools as a result of the spontaneous motor activity reverberating through a diffuse propriospinal network within the neonatal spinal cord ((Cazalets, 2005); Fig. 1 C, D). This observation was repeated in 20 out of 24 preparations.

To quantify the peak of injury-induced depolarization, high potassium (10 mM) was applied for 10 min before the impact to the same exemplar preparation (Fig. 1 A). KCl generated depolarizations that were smaller in VRrL5 (40.34%) and greater in VRrC2 (177.9%) compared to the ones induced by the following impact (Fig. 1 A).

A second exposure to 10 mM KCl after injury produced on both VRs the same depolarizations as the pre-impact application, demonstrating that the total number of functional motoneurons was unaffected by the impact in segments rostral and caudal to the lesion site (Fig. 1 A). Pooled data from five preparations showed that the peak of average injury-induced depolarizations from VRrL5 was significantly higher than the depolarizations elicited by 10 mM KCl (P < 0.001, Repeated measures analysis, n = 5; Fig. 1 E). Conversely, in the same group of preparations, the average injury-induced depolarization from VRrC2 was lower than the one elicited at lumbar levels (P < 0.001), and significantly lower than the depolarization determined by a second application of 10 mM KCl (Fig. 1 F; P = 0.046, Repeated measures analysis, n = 5). Notably, at both L5 and C2 levels, potentials elicited by rising KCl concentrations were comparable before and after the impact (Fig. 1 E, F). This confirms that an injury targeted to the low thoracic cord (T10) does not reduce the overall activation of motoneurons located in motor pools far from the injury site, which remain equally functional once directly activated by KCl. Furthermore, distinct lumbar segments of sham and injured spinal cords were treated with a selective marker for motoneurons in the ventral horns (SMI-32 antibody). Histological processing visualized a similar SMI-32 staining in the ventral cord of the sham and injured preparations, for both L1-L3 and L3-L5 segments (Fig. 1 H). Mean data from 49 slices from a total of eight animals (four sham intact and four injured spinal cords) confirmed no significant difference in the number of SMI-32 positive cells (P = 0.709, ANOVA), hence excluding the acute death of any lumbar motoneurons after the low thoracic injury and related spread depolarization.

Collectively, a physical insult to the mid-thoracic spinal cord triggers a transient and massive depolarization spreading along the entire spinal cord, suppressing the spontaneous motor activity that is derived synchronous among all neonatal VRs, yet without any cellular loss of lumbar motor pools.

### Calibrated impacts of increasing strength elicit higher peaks of depolarization

To assess the effects of varying degrees of severity of a spinal impact, increasing vertical displacements of the impactor rod were set. In each preparation, four different levels of compression (625 µm, 1250 µm, 1875 µm, and 2656 µm) were serially applied to the spinal cord at T10. A mild impact (625 µm) resulted in a moderate depolarization that was simultaneously recorded from VRrL2, VRrL5, and VRrL6, and which quickly recovered without perturbing the spontaneous baseline activity (Fig. 2 A). By increasing the strength of injury from 1250 to 1875 µm, progressively higher peaks of potentials were produced. At 1875 µm, the maximum level of depolarization was reached and could not be further increased even by the following most intense compression (2656 µm), likely due to the repetitive damage to the cord at the site of the impact (Fig. 2 A). Taking the VRrL5 recording of a sample experiment, the mildest trauma generated a depolarization of 1.4 mV, rising to 1.46 mV for 1250 µm, 3.52 mV for 1875 µm and eventually 2.41 mV during the strongest impact (2656 µm, Fig. 2 A). The impact at 2656 µm was used throughout the rest of the study to generate the most severe compression without completely transecting the neonatal spinal cord. Across all lumbar VRs, the extent of the depolarization elicited by 10 mM KCl was the same before and after the protocol of serial compressions, confirming that an injury at T10 does not affect the recruitment of motor pools below the lesion (Fig. 2 A). However, the spontaneous rhythmic motor activity arising synchronous from all VRs was largely reduced by the second impact (1250 µm) and up (Fig. 2 A). Pooled data from five experiments (Fig. 2 B) confirms that the amplitude of the injury-induced depolarization augments with stronger impacts, with significant higher peaks for VRrL5 (1875 Vs. 625 µm; P = 0.019, Kruskal-Wallis test), and for VRrL2 (1250 Vs 625 µm; P = 0.024, ANOVA). Moreover, for each injury intensity, depolarizations appeared by-and-by smaller, the farther the recording site was from the site of compression. This common trend is more evident for impacts at 1250 µm, where the peak of injury-induced depolarization was significantly higher for VRrL2 compared to VRrL6 (Fig. 2 B, P = 0.009; Kruskal-Wallis test, n = 5 for VRrL2 and VRrL5; n = 4 for VRrL6). Likewise, injury-induced depolarizations also appeared sooner in segments closer to the impact site, rather than from more caudal ones. Indeed, for all impact strengths, induced depolarization occurred first at L2, then at L5, and finally at L6 spinal segments, reaching a statistical significance for potentials elicited by the mildest impact (625 µm) at T10, between VRrL2 and VRrL6 (Fig. 2 C, P = 0.001, Kruskal-Wallis test, n = 5 for VRrL2 and VRrL5; n = 4 for VRrL6). Noteworthy, latency of depolarization recorded from each root was unchanged among the four intensities of injury showing that impact severity does not affect the velocity of depolarization spreading along the spinal cord.

The customized in vitro impactor allowed to consistently trace the features of injury-induced potentials for increasing severities of compression, showing that stronger impacts generate higher potentials without affecting their velocity of propagation from the impact site.

### Injury potentials propagate rostrally and caudally from the site of impact in ventro-dorsal directions

To better investigate the propagation of injury-induced depolarization along the entire spinal cord, we collected data from numerous VRs, out of a dataset of 44 preparations injured at the ventral aspect of T10 with the strongest impact (2656 µm tip displacement, Fig. 3 A). Injury potentials of different amplitude were recorded from distinct spinal segments, with the highest peaks from VRL1 and L2 being significantly larger than those derived at the extremities (Fig. 3 B, see Table 1 for statistical details). Injury potentials progressively slowed down the farther they were recorded from the impact site, with the lowest latency recorded at VRL1 (Fig. 3 C, see Table 2 for statistical details). Resulting velocity of the rostro-caudal conduction of injury-induced depolarizations from the site of impact to VRL1 (4.44 mm far from impact) was 0.03 ± 0.01 m/s, equal to the caudo-rostral conduction from the site of impact to VRT5 (4.83 mm far from impact, P = 0.451, Mann-Whitney test, n = 3 for T5 and n = 18 for L1).

**Figure 3.**
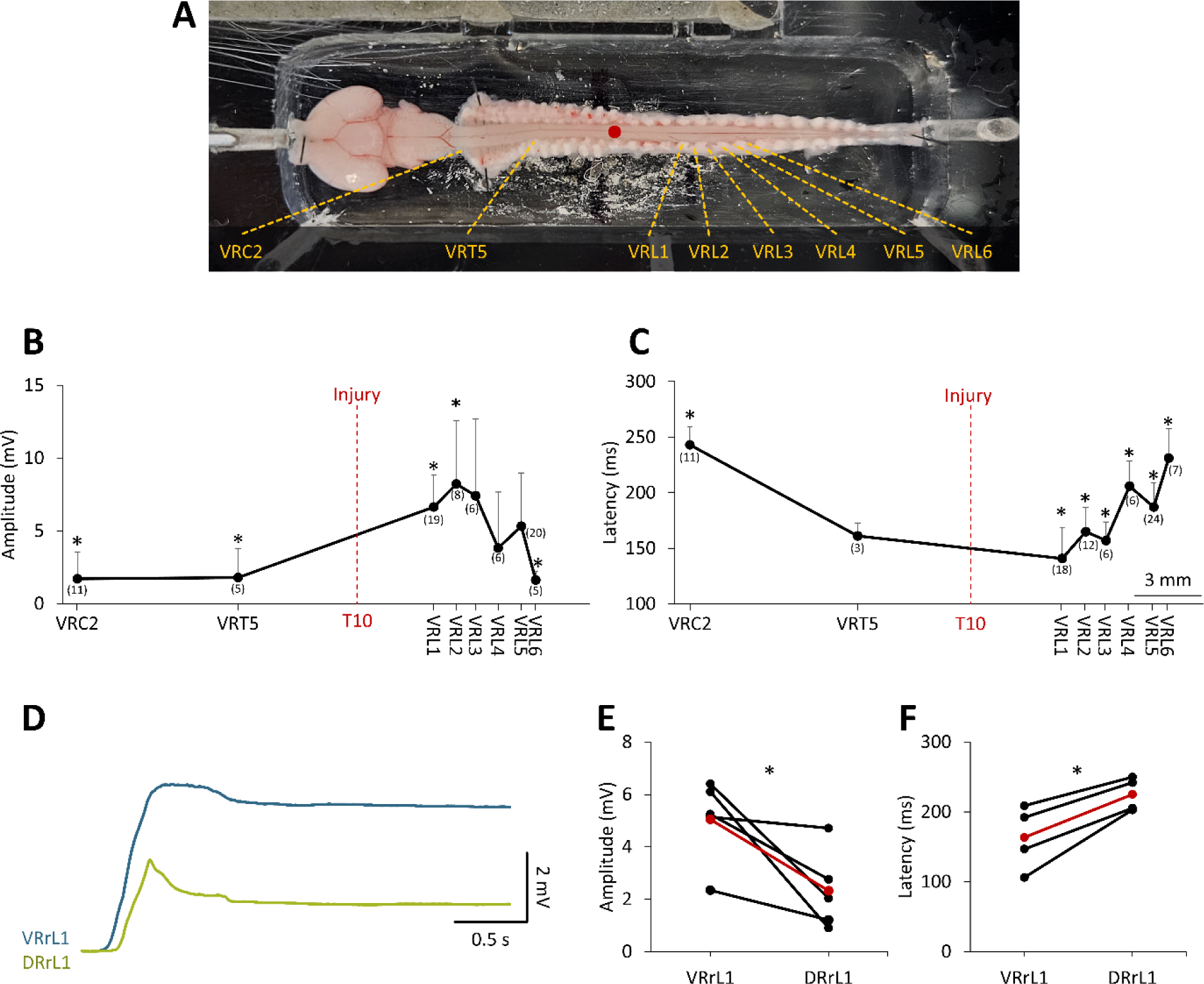
Impact-induced depolarization spreads from the injury site to the whole spinal cord. **A.** A ventral view of the CNS preparation with dorsal vertebrae attached. VRs recordings are taken from the VRs indicated by dotted yellow lines, while the injury site at the T10 segment is highlighted by a red dot. **B.** Mean amplitudes of injury potentials from several VRs. Red dotted line indicates the level of injury (T10). Number of experiments for each VR is indicated in brackets. Statistically significant amplitudes are indicated by *, as described in Table 1. **C.** Mean latencies of injury potentials from several VRs. Red dotted line indicates the level of injury (T10). Number of experiments for each VR is indicated in brackets. Statistically significant amplitudes are indicated by *, as described in Table 2. **D.** Superimposed mean traces from simultaneous recordings of injury potentials from both, DR (green trace) and VR (blue trace), at L1 (n = 4). **E, F.** Injury potentials from DRrL1 are significantly smaller (E; *P = 0.041) and slower (F; *P = 0.015) than recorded from VRrL1. Red dots and line show average values.

**Table 1.** Amplitude values of impact-induced depolarizations from different VRs. P values correspond to Kruskal-Wallis test.

| Amplitude |  |  |  |  |  |  |  |  |
| --- | --- | --- | --- | --- | --- | --- | --- | --- |
| Vs | VRC2 | VRT5 | VRL1 | VRL2 | VRL3 | VRL4 | VRL5 | VRL6 |
| VRC2 |  | P>0.05 | P<0.01 | P<0.01 | P>0.05 | P>0.05 | P>0.05 | P>0.05 |
| VRT5 | ns |  | P>0.05 | P<0.05 | P>0.05 | P>0.05 | P>0.05 | P>0.05 |
| VRL1 | * | ns |  | P>0.05 | P>0.05 | P>0.05 | P>0.05 | P<0.05 |
| VRL2 | * | * | ns |  | P>0.05 | P>0.05 | P>0.05 | P<0.05 |
| VRL3 | ns | ns | ns | ns |  | P>0.05 | P>0.05 | P>0.05 |
| VRL4 | ns | ns | ns | ns | ns |  | P>0.05 | P>0.05 |
| VRL5 | ns | ns | ns | ns | ns | ns |  | P>0.05 |
| VRL6 | ns | ns | * | * | ns | ns | ns |  |
| Average | 1.72 | 1.79 | 6.64 | 8.23 | 7.42 | 3.83 | 5.33 | 1.6 |
| SD | 1.84 | 1.99 | 2.20 | 4.37 | 5.28 | 3.85 | 3.67 | 0.59 |

**Table 2.** Latency values of impact-induced depolarizations from different VRs. P values correspond to Kruskal-Wallis test.

| Latency |  |  |  |  |  |  |  |  |
| --- | --- | --- | --- | --- | --- | --- | --- | --- |
| Vs | VRC2 | VRT5 | VRL1 | VRL2 | VRL3 | VRL4 | VRL5 | VRL6 |
| VRC2 |  | P>0.05 | P<0.001 | P<0.001 | P<0.001 | P>0.05 | P<0.05 | P>0.05 |
| VRT5 | ns |  | P>0.05 | P>0.05 | P>0.05 | P>0.05 | P>0.05 | P>0.05 |
| VRL1 | * | ns |  | P>0.05 | P>0.05 | P<0.01 | P<0.01 | P<0.001 |
| VRL2 | * | ns | ns |  | P>0.05 | P>0.05 | P>0.05 | P<0.05 |
| VRL3 | * | ns | ns | ns |  | P>0.05 | P>0.05 | P<0.05 |
| VRL4 | ns | ns | * | ns | ns |  | P>0.05 | P>0.05 |
| VRL5 | * | ns | * | ns | ns | ns |  | P>0.05 |
| VRL6 | ns | ns | * | * | * | ns | ns |  |
| Average | 243.13 | 161.2 | 140.81 | 165.03 | 157.1 | 205.97 | 187.19 | 231.31 |
| SD | 16.36 | 11.79 | 28 | 22.14 | 16.7 | 22.84 | 21.68 | 26.62 |

To gain insights on the dorsal-ventral propagation of injury-induced depolarization, we simultaneously derived from both VRrL1 and DRrL1 while impacting the ventral side of the cord at T10. Data pooled from many experiments (Fig. 3 D) indicates that the impact leads to injury potentials that propagate also to the dorsal part of the cord, although they appear smaller (P = 0.041, paired t-test, n = 5) and spread more slowly (P = 0.015, paired t-test, n = 4) than ventrally elicited potentials.

Present data indicates that a physical impact to the spinal cord elicits a strong wave of depolarization that departs from the site of injury and invests the entire spinal cord with the same velocity, affecting also dorsal segments. This observation provides the rationale for ascertaining the functionality of spinal networks above and below the site of injury.

### An impact generates potentials that equally propagate to both sides of the cord, and disconnects the lumbar cord from descending respiratory input

To confirm the symmetrical propagation of injury-induced depolarizations along both sides of the cord, simultaneous VR recordings were obtained from both left and right VRs at L1, in response to a physical impact at T10. In a sample experiment, continuous recordings were acquired from VRlL1, VRrL1, and VRrC2 (Fig. 4 A). After the impact, VR injury-induced potentials peaked at 10.15 mV and 11.38 mV for left and right VRs, respectively. Average data from four experiments indicated an equal extent of impact-induced depolarizations on both sides of the L1 spinal segment (Fig. 4 B, P > 0.999, Wilcoxon matched-pairs signed-ranks test).

**Figure 4.**
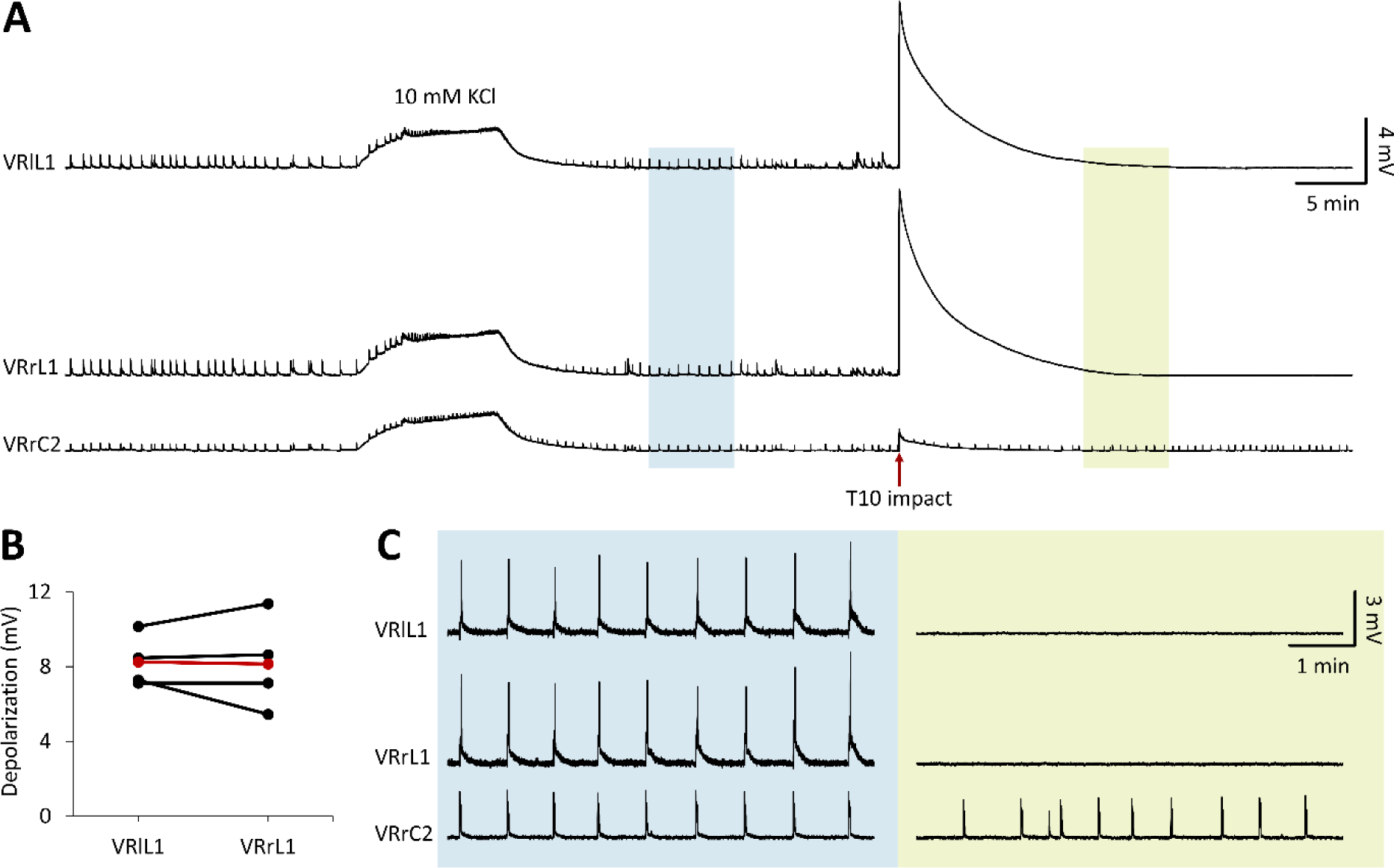
Impact evokes equal bilateral injury potentials and disconnects lumbar motor pools from descending respiratory input. **A.** Continuous and simultaneous recordings from VRrL1, VRlL1, and VRrC2 showing the exposure to a high potassium solution (10 mM, 10 min) and to the following impact at T10. **B.** The plot visualizes the equal amplitude of injury-induced depolarizations recorded from left and right L1 VRs (n = 4). Red dots and red line correspond to average values. **C.** Magnifications correspond to the pale regions of continuous traces in A, and highlight rhythmic respiratory bursts in control (blue panel) and 21.8 min after the impact (green panel). Fictive respiration originating from brainstem structures is maintained at VRrC2 but disappeared from lumbar VRs due to the interruption of descending input beyond the site of impact.

Furthermore, in the same sample experiment, spontaneous rhythmic bursts (0.02 ± 0.01 Hz) originating from respiratory networks in the brainstem (Mohammadshirazi et al., 2023; Apicella and Taccola, 2023) were simultaneously recorded in control from cervical and lumbar VRs (Fig. 4 C, left). In injured preparations, fictive respiration disappeared from all lumbar VRs, while after 20 mins from the impact, spontaneous rhythmic bursts from VRC2 persisted with a frequency similar to control (0.02 ± 0.01 Hz, Fig. 4 C, right). This observation was repeated in seven preparations, confirming both the injury-induced suppression of lumbar respiratory events, and the endurance of fictive respiration from cervical VRs with unchanged frequency from pre-injury controls (0.05 ± 0.03 Hz from 20 min pre-injury, 0.05 ± 0.02 Hz from 20 min post-injury, P = 0.709, paired t-test).

In summary, the equal magnitude of bilateral injury potentials propagating to lumbar VRs confirms the midline location of the impact. Moreover, the disappearance of respiratory bursts below the site of injury indicates that lumbar motor pools are completely disconnected from supraspinal respiratory centers.

### Impact causes extensive neuronal loss at the contusion site and completely disconnects ascending afferent input

Disappearance of respiratory episodes from the lumbar cord indicates that descending respiratory input from the brainstem are blocked at the level of impact. To investigate whether also the conduction of ascending input is interrupted by the impact, we recorded ascending input from VRs, as evoked by continuous electric stimulations (intensity = 100 µA, pulse duration = 0.1 ms, frequency = 0.1 Hz) of sacrocaudal afferents (Etlin et al., 2010). Simultaneous recordings were taken above and below the level of impact. In a sample experiment, single reflex responses in control were 1.26 and 0.13 mV as recorded from VRrL5 and VRrC2, respectively (blue traces in Fig. 5 A). At the peak of injury-induced depolarization, both responses vanished (Fig. 5 A). After 38 s from the impact, reflex responses from VRrL5 reappeared and eventually stabilized after 8 min, albeit reduced in amplitude to 41 % of pre-impact control. Cervical responses were completely abolished (green traces in Fig. 5 A). The disappearance of cervical reflexes after the impact was replicated in nine out of nine preparations.

**Figure 5.**
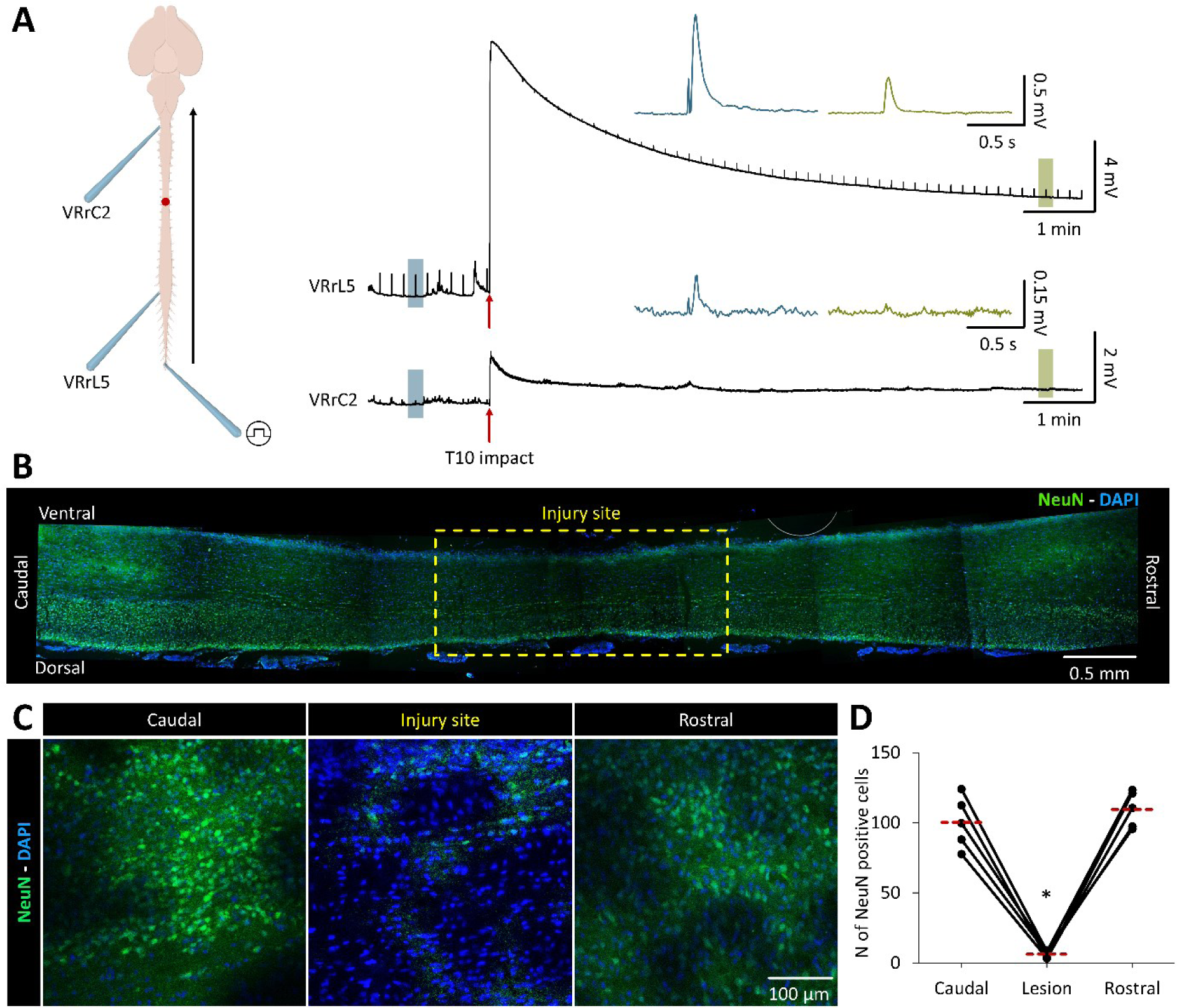
Contusion suppresses ascending conduction of afferent input and causes massive neuronal death at the site of ventral impact. **A.** The cartoon depicts the CNS preparation with the impact site on the ventral aspect of T10 (red dot). Extracellular electrodes are positioned at C2 and L5 rVRs, and repetitive electrical pulses (0.1 Hz, 100 µA, duration = 100 µs) are supplied to cauda equine to elicit ascending input (arrow). Right traces show simultaneous recordings from VRrL5 and VRrC2 with reflex responses appearing in control and magnified in the blue insert. After the depolarization induced by the impact (red arrow), evoked motor responses are abolished on both VRs. During repolarization, responses progressively reappear on lumbar VR, while lumbar reflexes become visible again after 38 s from the impact and recover towards the original size by the time (8 min, top green insert). Contrariwise, reflexes from VRrC2 do not recover (bottom green insert). **B.** Reconstruction of sagittal slices of a spinal cord (caudal left, rostral right, ventral up, dorsal down) as processed with DAPI and NeuN staining. A massive cellular loss is visible on the ventral aspect of the impact site. The base of the dotted yellow rectangle is calibrated to the width of the impactor tip. C. Magnifications of horizontal slices stained with DAPI and NeuN, and collected from serial spinal segments at caudal level (T11, left), injury site (T10, middle) and rostral spinal cord (T9, right The lack of NeuN (green) staining at the site of impact indicates extensive neuronal loss. D. The plot quantifies the statistical reduction of NeuN-positive cells at the injury site compared to both rostral and caudal segments (*P < 0.001). Red dotted lines correspond to the average number of NeuN positive cells.

To exclude that the reduced lumbar reflex amplitude arose from an interference produced by the impactor movement, rather than from a real depolarization caused by the injured tissue, in a subset of experiments, lumbar responses were allowed to recover after being transiently abolished by a first impact at T10. Then, the spinal cord was completely transected at L1 level (Sl. Fig. 3 A, B) and a second impact at T10 was performed, which did not evoke any injury potentials from the disconnected caudal cord nor varied the amplitude of reflex responses (Sl. Fig. 3 B). Noteworthy, the second impact still elicited an injury potential from the rostral cord (Sl. Fig. 3 B).

To visualize the anatomical damage caused by the impact, histological assessments were performed on sagittal sections of the entire spinal cord. The ventral spinal cord at the site of impact (dotted yellow rectangle) showed negligible neuronal labeling for NeuN due to an extensive cell loss (Fig. 5 B).

In another example, magnifications of horizontal slices from serial close spinal segments confirmed a lower number of NeuN positive cells at the injury site from 5 injured spinal cords (Fig. 5 C). Pooled data from five experiments demonstrated the significant reduction of NeuN-positive cells at the injury site (T10) compared to rostral (T9) and caudal (T11) segments (P < 0.001, ANOVA; see Fig. 5 D).

This histological evidence describes a massive neuronal damage at the site of injury and corroborates the functional deficits reported above, namely the complete interruption of longitudinal spinal input at the level of impact.

### Cord oxygenation drops after a spinal impact

After an SCI, systemic hypotension and pericyte constriction of spinal capillaries decrease spinal oxygen delivery, reducing oxygen concentration on spinal tissues (Partida et al., 2016; Li et al., 2017). To quantify PO2 in spinal cord tissue during contusion, an oximeter sensor was positioned 100 µm deep into the cord on the anterior funiculus between L1 and L2 VRs, while continuous electrophysiological signals were derived from VRrL1. Impact at T10 (red arrow) induced a large depolarization (5.23 mV, Fig. 6 A), which recovered to baseline after 12 min. In a sample experiment, tissue oxygen was continuously monitored, showing PO2 values oscillating between 20.44 and 50.91 Torr in control (Fig. 6 B). Immediately after the impact (red arrow), PO2 dropped to 8.12 Torr, but recovered pre-impact values after 10 min, perfectly matching the profile of DC level changes (Fig. 6 A, B). The time course of average PO2 from nine preparations indicated that tissue oxygen content in control (31.19 ± 7.36 Torr) dropped to 11.68 ± 4.03 Torr after the impact, and then slowly recovered to the 78.74 % of control after 30 min (Fig. 6 C).

**Figure 6.**
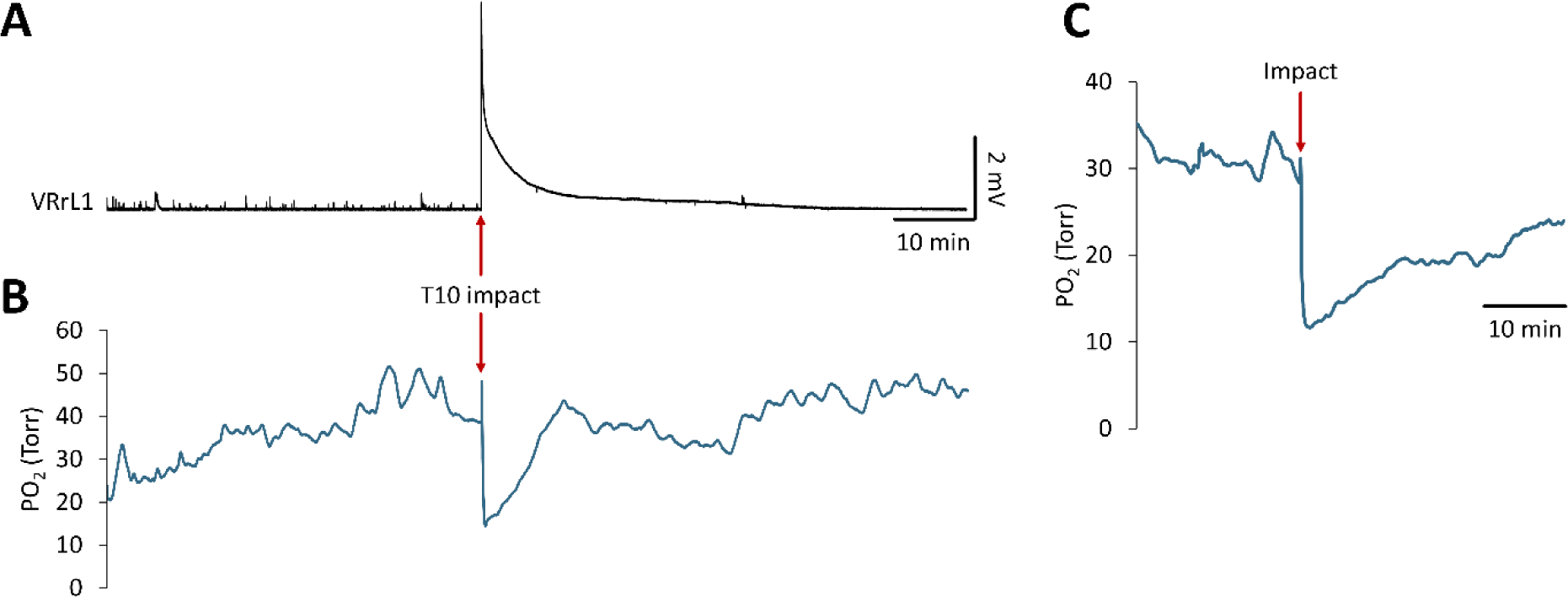
Impact drops oxygen tension of the spinal cord with a pattern that resembles the profile of injury-induced depolarization. **A.** Continuous trace from VRrL1 with a large depolarization at the site of impact at T10 (red arrow). **B.** Simultaneous PO2 measurements performed from the anterior funiculus between L1/L2 VRs in the same experiment as in A. PO2 drops right after the impact, eventually recovering to baseline, mirroring the depolarization profile in A. **C.** Average spinal PO2 profile before and after the impact (red arrow, n = 9).

Oxygen consumption for in vitro preparations parallels the level of cellular activity (Wilson et al., 2003). To provide a reference for spinal oxygen consumption during a large depolarization, the CNS was perfused for ten minutes with a modified Krebs solution containing 10 mM KCl. High K^+^ (10 mM) induced a mean depolarization of 1.83 ± 0.54 mV from VRL1, while average PO2 measured from the L1 spinal segment dropped to 9.54 ± 2.14 Torr (Sl. Fig. 4 A).

The link between the increased neural activity induced by a large depolarization and the PO2 consumption was confirmed using a CNS preparation that underwent a functional inactivation through heat-shock (100°C) and then a continuous perfusion with oxygenated Krebs. Here, no depolarization was recorded from VRrL5 after exposure to high K^+^ (10 mM), while the oximeter probe inserted at L1 spinal level derived a mean PO2 of 528 ± 8.74 Torr equal to pre-K^+^ control values. In the same preparation, the spinal impact did not elicit any depolarizations from VRrL5, with PO2 measurements that remained unchanged before and during the impact (505.76 ± 2.57 in control and 508.75 ± 3.16 Torr during impact, Sl. Fig. 4 C).

Collectively, the impact-induced drop in PO2 parallels the kinetics of impact-induced depolarization. Furthermore, a spinal impact largely reduced spinal tissue oxygen to levels comparable to a strong network activation using 10 mM K^+^.

### Impact transiently suppresses lumbar motor reflexes

A compression of the spinal cord is followed by a spinal shock, characterized by the suppression of motor evoked responses lasting beyond the moment of the first insult (Ditunno et al., 2004). To confirm the presence of a shock phase in our in vitro SCI model, stimuli were continuously supplied to sacrocaudal afferents (frequency = 0.1 Hz; intensity = 100 µA, 5 × Th; pulse duration = 0.1 ms) while motor reflexes were derived from VRrL5 in control and after the impact (Fig. 7 A). Firstly, the concentration of potassium was raised to 10 mM, eliciting a depolarization of 4.05 mV at steady state. At the peak of the K^+^-induced depolarization, reflexes were suppressed, but recovered to baseline during the subsequent washout in normal Krebs (Fig. 7 A). After the washout, motor evoked responses appeared transiently suppressed at the peak of the impact-induced depolarization (9.69 mV) but, as the baseline repolarized, also pre-impact values fully recovered after 31.06 min from the impact (Fig. 7 A). A second exposure to high K^+^ concentrations (10 mM) evoked a depolarization of 4.55 mV, which abolished motor reflexes until a normal Krebs solution was perfused and led to a full recovery of reflexes (Fig. 7 A). The profile representing changes in the amplitude of reflex responses throughout the experiment displays a complete suppression of motor reflexes in correspondence to a spinal cord depolarization of about 4 mV, regardless of whether it was elicited by perfusing the whole preparation with high potassium ions or by applying a localized impact at T10 (red arrow, Fig. 7 B). Similar evidence was obtained from five preparations, where trains of pulses (frequency = 0.1 Hz, intensity = 1.6-6.15 Th, pulse duration = 0.1ms) applied to sacrocaudal afferents evoked spinal reflexes from VRrL5 (peak amplitude = 0.77 ± 0.2 mV). After 27.91 ± 6.06 s from the impact at T10, electrically evoked responses reappeared, and recovered to 90% of pre-impact values after 18.25 ± 12.2 minutes. In the same five preparations, the time of reappearance of the first reflex after impact was not correlated to the amplitude of the injury potential (correlation coefficient = −443, P = 0.455). Through multiple simultaneous recordings, comparable transient suppressions of DRVRPs were reported across VRs at spinal segments L1, L2, L4, L5, L6 on both sides.

**Figure 7.**
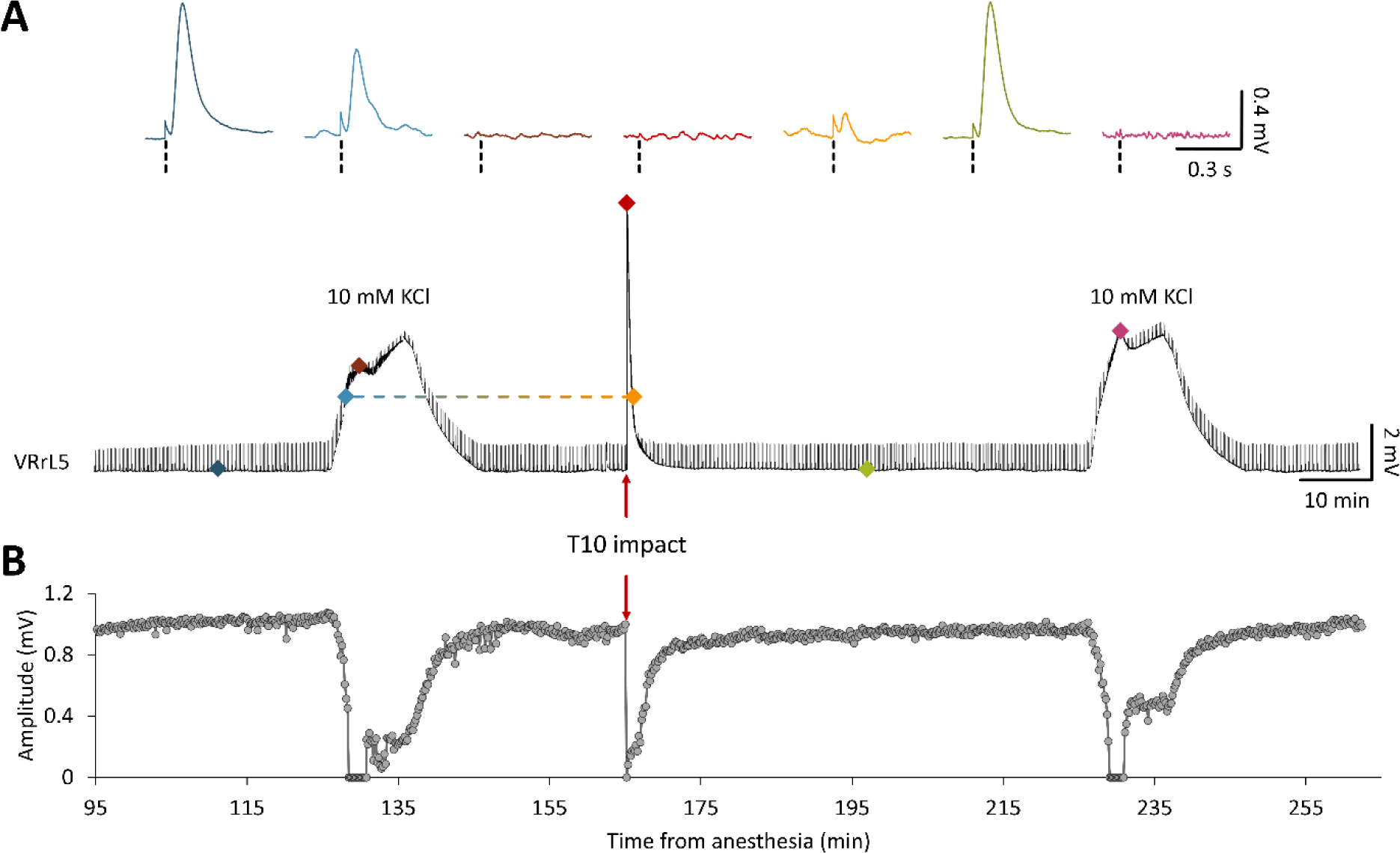
Motor reflexes vanish at the peak of both, chemically- and impact-induced depolarizations. **A.** A 178 min long recording from VRrL5 during the continuous delivery of electrical pulses to sacrocaudal afferents (0.1 Hz) to evoke motor responses. Motor reflexes are traced in control, during high K^+^ (10 mM) perfusion, wash out, impact to T10 (red arrow) and a second K^+^ application. Top inserts magnify single reflexes (dotted vertical lines correspond to artifacts of stimulation) for distinct instants of the experiment as indicated by the colored dots below. At the top of each depolarization, motor reflexes are suppressed (brown, red and purple top traces). **B.** Time course of reflex amplitude for the trace in A demonstrates that reflexes vanish (amplitude = 0 mV) at the peak of depolarizations, regardless of whether they are elicited by the perfusion of the entire CNS with high K^+^ or by a localized impact to the cord.

In summary, in the current study, the calibrated and localized impact to the cord has always been followed by a transient suppression of evoked reflexes from spinal motor pools.

### A thoracic impact alters electrically-induced fictive locomotor patterns

Results collected so far indicate that, after an impact, the entire spinal cord experiences a transient large depolarization, with neuronal death only at the injury site. To explore whether the depolarization induced by the impact affects the functionality of lumbar spinal networks for locomotion, stereotyped trains of rectangular pulses (frequency = 2 Hz, intensity = 1-5 × Th, pulse duration = 0.1 ms) were applied to sacrocaudal afferents for 80 seconds. In response to stimulation, episodes of locomotor-like oscillations alternating between flexor and extensor commands, and between left and right motor pools were recorded from both VRL2 and from one VRL5, in control and at different time points after the impact (15, 60 and 120 min post-SCI). In a sample experiment, fictive locomotor patterns recorded in control from VRrL2 were characterized by a cumulative depolarization of 0.7 mV with 28 superimposed alternating cycles (CCFhomolateral = −0.70, CCFhomosegmental = −0.87), defined by a peak amplitude of 0.33 ± 0.08 µV and a period of 2.89 ± 0.74 s (Fig. 8 A and magnification at steady state in Fig. 8 B). In the same preparation, the impact reduced cumulative depolarization (0.5 mV, 15 min post-SCI), generating smaller (0.16 ± 0.06 µV, 15 min post-SCI) and slightly less regular locomotor-like oscillations (period CV = 0.28, 15 min post-SCI Vs. period CV in control = 0.26), regardless of their unchanged number (28, 15 min post-SCI). Although some features of fictive locomotion eventually recovered to control values, cumulative depolarization, cycle amplitude and periodicity were still reduced even after two hours from the impact (Fig. 8 A and magnified at steady state in Fig. 8 B).

**Figure 8.**
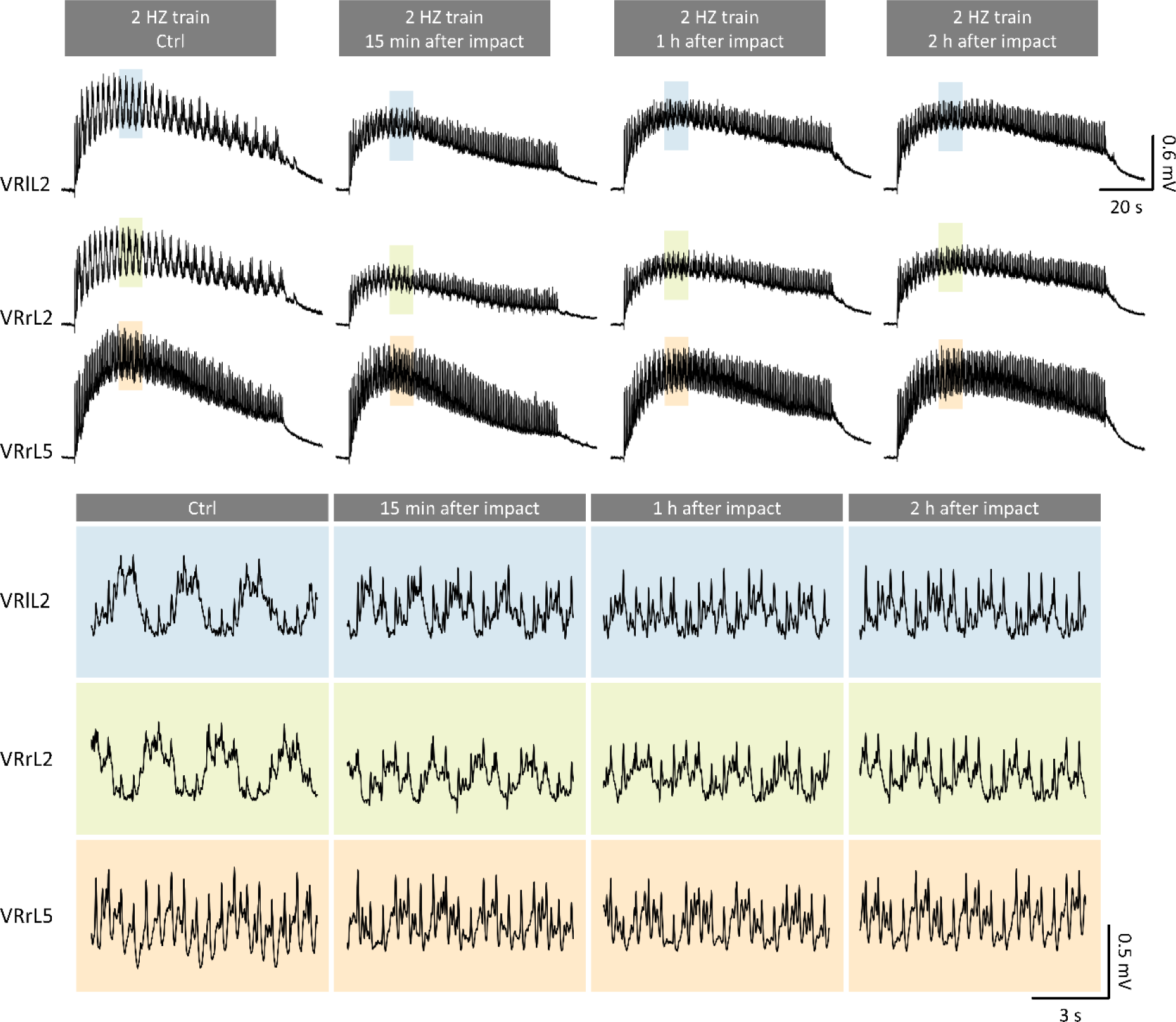
Electrically-induced fictive locomotion is affected by a localized thoracic compression. **A.** Serial 2 Hz trains of stereotyped rectangular pulses (intensity = 125 µA, duration = 0.1 ms) are applied to sacrocaudal afferents to evoke epochs of locomotor-like oscillations from VRlL2, VRrL2, and VRrL5. Fictive locomotion patterns were recorded in control, and then 15 minutes, one hour, and two hours after injury. **B.** Magnifications of simultaneous traces (blue for VRlL2, green for VRrL2, and orange for VRrL5) correspond to oscillations captured at steady state in A (shaded rectangles). Note the out-of-phase cycles recorded from the three VRs, with reduced amplitude and periodicity after the impact.

Pooled data from seven experiments confirms that the impact unaltered several characteristics of fictive locomotion (Sl. Fig. 5) but did reduce cumulative depolarization (Fig. 9 A; n = 6, P = 0.002, repeated measures analysis) and amplitude of cycles from VRrL2 (Fig. 9 B; n = 6, P < 0.001, repeated measures analysis). In addition, duration of fictive locomotion episodes from VRrL2 after 60 minutes from the impact (Fig. 9 C; n = 7, P = 0.031, repeated measures analysis), and period of cycles of VRrL2 after 15 and 60 minutes from the impact, were significantly lower than in control (Fig. 9 D; n = 7, P = 0.008, repeated measures analysis). Similarly, 15- and 60-min post-impact, episodes from VRrL5 were faster than in the control group (Fig. 9 E; n = 6, P = 0.002, Friedman test), as well as more irregular at 15 minutes post-impact (Fig. 9 F; n = 7, P = 0.006, repeated measures analysis). Notably, after injury, oscillations from both extensor and flexor commands (Fig. 9 G; homolateral CCF, n = 7, P = 0.013, repeated measures analysis), as well as from the left and right sides of the cord (Fig. 9 H; homosegmental CCF, n = 7, P = 0.001, repeated measures analysis) exhibited poorer alternating coupling than controls.

**Figure 9.**
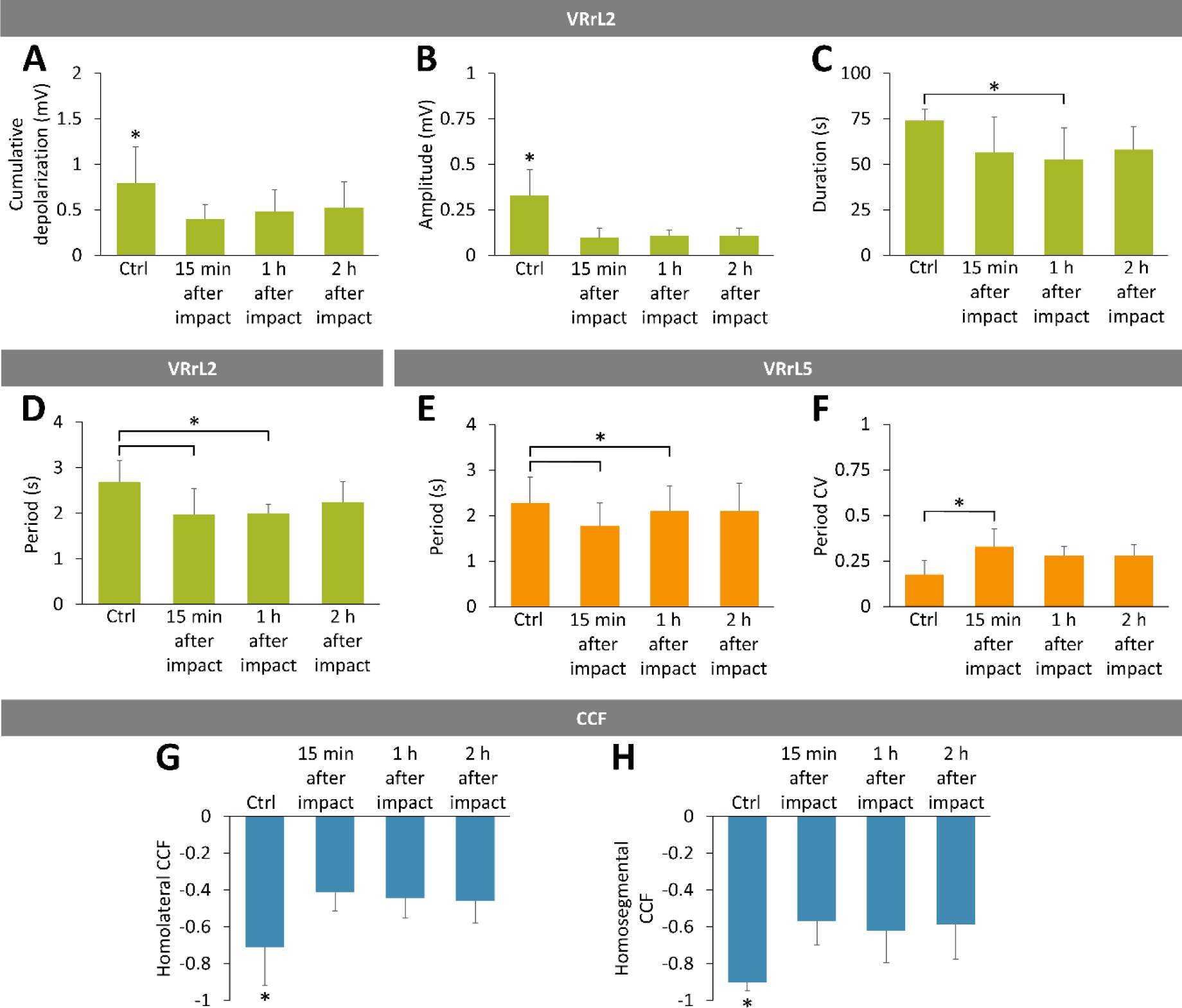
Impact perturbs the features of electrically-induced fictive locomotion. **A-D.** Green bars describe average values for the main descriptors of fictive locomotor patterns reported from VRrL2 in control and at 15 minutes, 1 hour, and 2 hours following the injury. **A.** Cumulative depolarization significantly decreases after impact (*P = 0.002). **B.** Impact largely reduces the amplitude of oscillations (*P < 0.001). **C.** Episodes of fictive locomotion are shorter one-hour after the impact (*P = 0.031). **D.** Period of oscillations is significantly smaller 15 minutes and one-hour post-impact (*P = 0.008). **E-F.** Orange bars describe average values for the main descriptors of fictive locomotor patterns reported from VRrL5 in control and at 15 minutes, 1 hour, and 2 hours following the injury. **E**. Periods of FL oscillations 15 minutes and one hour after injury are significantly shorter than in the control group (*P = 0.002). **F.** Period CV is higher than in control only at 15-minute post-injury (*P = 0.006). **G.** Phase coupling between extensor and flexor commands (homolateral CCF, *P = 0.013) is poorer after the impact. **H.** Phase coupling between the left and right output (homosegmental CCF, *P = 0.001) reduces post injury.

In summary, a calibrated impact to the thoracic cord affects the functionality of lumbar locomotor circuits, generating less coordinated locomotor-like oscillations, with shorter and faster cycles of locomotor-like patterns especially from flexor motor pools.

### Impact-induced depolarization is sustained by chloride ions

To investigate whether ionic disbalances sustain the depolarization that follows the impact, separate experiments considered injuring the cord during perfusion with either of the three modified Krebs solutions containing low concentrations of chloride (Cl^−^), calcium (Ca^2+^), and potassium (K^+^) ions, respectively. Continuous recordings were performed from preparations initially perfused with normal oxygenated Krebs solution, and then with one of the low-ion Krebs solutions for 30-90 min before the impact and for 15 min afterwards. As soon as a single low-ion solution was applied, the DC level of the baseline recorded from VRrL5 hyperpolarized and, after 18 min, reached a steady-state mean level of −10.42 ± 2.23 mV for low Cl^−^ (n = 5), −0.49 ± 0.39 mV (n = 7) for low Ca^2+^ and −1.11 ± 0.75 mV for low K^+^ (n = 7; Sl. Fig. 6 A).

Whether low-ion solutions affected spinal synaptic transmission was verified by continuously monitoring the reflex responses elicited from VRrL5 in response to trains of weak electric pulses (frequency = 0.1 Hz, intensity = 50-160 µA, 2-8 × Th) applied to sacrocaudal afferents. Three pairs of superimposed sample traces from three preparations show that reflex amplitudes were differently affected by the transition from a normal Krebs solution (blue traces) to each low-ion perfusion (green traces). In particular, compared to the normal Krebs solution, the peak of responses remained unchanged when perfusing low Cl^−^ (Fig. 10 A, left), while it reduced to 30.69 % during perfusion with low Ca^2+^ solution (Fig. 10 A, middle) and augmented to 117.88 % after the transition to low K^+^ (Fig. 10 A, right). Pooled data from many experiments confirms that the peak of mean reflexes was unchanged by low Cl^−^ (n = 6, P = 0.923, paired t-test), while it significantly reduced after low Ca^2+^ (n = 7, P = 0.001, paired t-test) and it increased with low K^+^ (n = 7, P = 0.017, paired t-test; Fig. 10 B). Conversely, latency of responses was only affected by the transition to the low Cl^−^ solution (P = 0.001, paired t-test; Fig. 10 C, left) without any changes appearing with low Ca^2+^ (P = 0.069, paired t-test, Fig. 10 C, middle) or low K^+^ (P = 0.297, paired t-test; Fig. 10 C, right). Impacts occurring during perfusion with low-ion solutions generated different peaks and profiles of injury-induced potentials. Comparison between three mean traces recorded for up to 3.5 min after the impact (red arrows) demonstrates that low Cl^−^ concentrations (n = 6, Fig. 10 D, left) generate higher peaks of injury potentials compared to the other two modified Krebs solutions. Furthermore, despite a lower peak of depolarization, low Ca^2+^ broadened the average injury potentials with the appearance of a delayed component in the repolarizing phase (n = 7; Fig. 10 D, middle). Low K^+^ perfusion showed a peak similar to low Ca^2+^ depolarizations, but with a sharper repolarizing phase (n = 7; Fig. 10 D, right).

**Figure 10.**
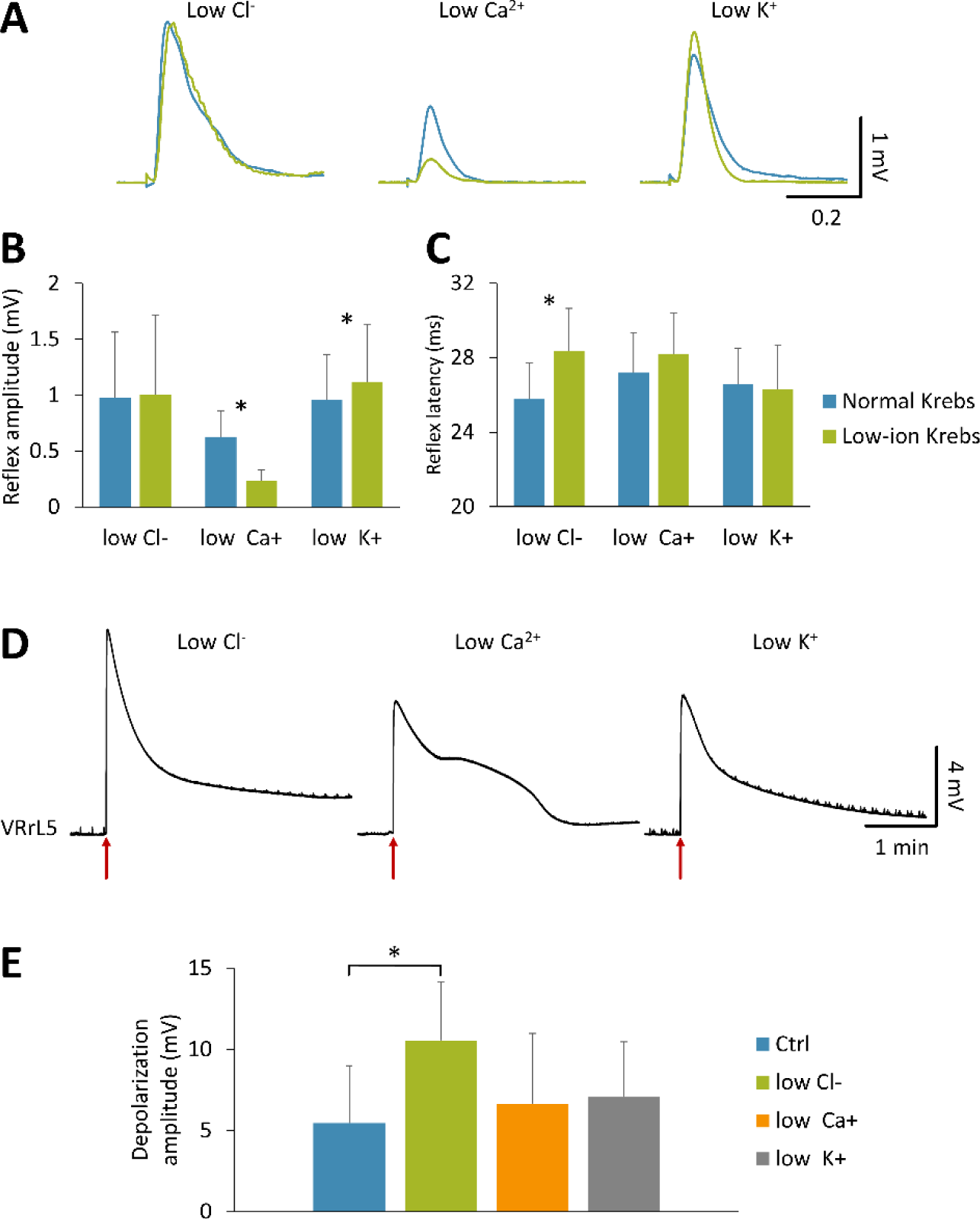
Low-ion solutions differently influence synaptic transmission and impact-induced depolarization. **A.** Superimposed pairs of electrically-induced reflex responses representing transition from normal Krebs solution (blue traces) to modified Krebs solutions (green traces) with low Cl^−^ (left), low Ca2^+^ (middle), and low K^+^ (right) concentrations. **B.** Mean amplitude of reflex responses in normal Krebs (blue bars) or low-ion solutions (green bars) point out a significantly smaller peak during application of low Ca^2+^ (P < 0.001), and a higher amplitude after transition to low K^+^ (P = 0.017) solutions. **C.** Mean values of latency of reflex responses in normal Krebs (blue bars) and low-ion solutions (green bars) report significantly slower responses when the low Cl^−^ solution is perfused (P = 0.001). **D.** Four-minute traces from VRrL5 related to the average profiles of injury potentials during perfusion with low-ion solutions (left, low Cl^−^, n = 6; middle, low Ca^2+^, n = 7; right, low K^+^, n = 7). **E.** Average peak amplitude of impact-induced depolarizations during perfusion in control Krebs and low-ion solutions. Low Cl^−^ solution augments the amplitude of depolarization (*P < 0.05).

Comparison among the mean amplitude of injury potentials generated by the impact during perfusion in normal Krebs (5.46 ± 3.54 mV; n = 23) and in the presence of the three low-ion solutions indicated a significantly higher depolarization for impacts occurring in low Cl^−^ (10.56 ± 3.57 mV, P = 0.048, Kruskal-Wallis test, n = 6, Fig. 10 E). Nevertheless, after impact in low Cl^−^, reflex responses were suppressed with a time course reminiscent of post-injury reflexes in normal Krebs solution, with a first reappearance of responses after 20.64 ± 6.15 s from the impact and the recovery to 90% of pre-impact values after 11.08 ± 4.61 mins.

Impacts in the presence of the modified Krebs solutions revealed the distinct role of Cl^−^ ions in sustaining the extent of injury potentials, albeit the duration of spinal shock and the suppression of reflex responses were comparable among the different media.

### An impact to the spinal cord alters cortical glial

To evaluate the impact of spinal injury on brain structures, the cerebral cortex was examined at two time points: the acute phase (25 minutes post-injury) and late phase (two hours post-injury). Given that the experiments were conducted during the peak of astrogenesis in rats, the density of astrocytes in the cerebral cortex was assessed as an indicator of the potential effects of the spinal insult. Astrocytes were identified through immunostaining of cortical samples for the S100b marker, followed by counterstaining with DAPI. The density of astrocytes was calculated by dividing the number of S100b-positive cells by the total cell count. In the dorsomedial cortex (at the level of the primary motor area, M1, Fig. 11 A), average astrocyte density was significantly reduced 25 minutes post-injury (25.85 % of sham) with a partial recovery two-hours post-injury (55.78 % of sham; Fig. 11 B, D; P = 0.033, Kruskal-Wallis test, n = 3, 3, 6). Contrarywise, in the ventrolateral cortex (including both primary and secondary somatosensory areas, S1 and S2, Fig. 11 A), astrocyte mean density was 54.35 % of sham, 25 min post-injury, and then significantly decreased two-hours post-injury (50.90 % of sham; Fig. 11 C, E; P = 0.037, Kruskal-Wallis test, n = 3, 4, 7). In summary, density of cortical astrocytes transiently changed, first at the level of the primary motor area and subsequently in both somatosensory areas, mirroring the spreading of depolarization from the injury site in the spinal cord.

**Figure 11.**
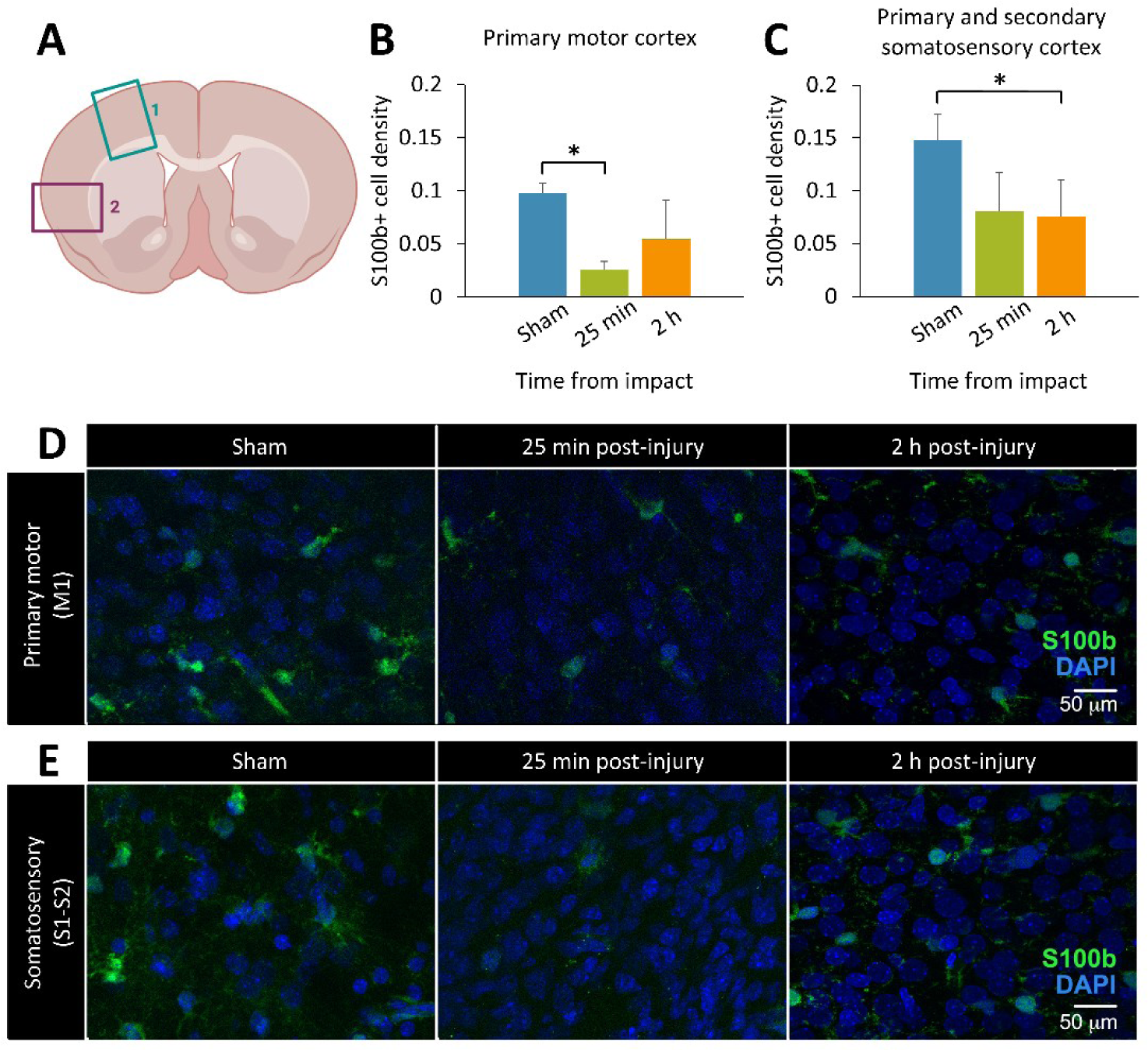
Remote changes in cortical glia occur after spinal damage. **A.** Illustration showing a coronal section of the cerebral cortex used for analysis. Square 1 indicates the dorsomedial cortex, including the primary motor area; Square 2 indicates dorsolateral and ventrolateral cortices, including both primary and secondary somatosensory cortices. **B.** Average S100b+ cell density in the primary motor cortex, 25 minutes and 2 hours post-injury, compared to sham. **C.** Average S100b+ cell density in the primary and secondary somatosensory cortices, 25 minutes and 2 hours post-injury, compared to sham. **D.** Representative confocal images from the primary motor cortex in sham, 25 minutes and 2 hours post-injury. S100b (in green) was used to label astrocytes, DAPI (in blue) was used to counterstain total cell nuclei. Images were taken at 60X oil objective magnification. **E.** Confocal images from the somatosensory cortex at the same time points as in D.

## Discussion

The current study is centered around tracing the immediate consequences of a traumatic injury to the spinal cord. For the purpose, we designed a novel experimental platform composed of a classical electrophysiological set up for multiple and simultaneous nerve recordings and stimulation, combined with an oximetric implantable probe, and associated with the invention and use of an ad hoc low-noise mini-impactor. This experimental infrastructure has been tailored to best exploit the innovative version of the whole CNS isolated from neonatal rats we recently introduced (Mohammadshirazi et al., 2023). To provide a more physiological and stable site for a traumatic impact, we adopted a more conservative surgery that maintained dorsal vertebral laminae and DRGs mostly intact. The impactor was carefully shielded to simultaneously allow VR recordings, and was created as to allow consistent and calibrated focal impacts through a fully programmable software interface. Each impact triggered a transient and massive depolarization spreading from the injury site to the whole spinal cord, symmetrically propagating across the left and right sides of the cord, and also to the dorsal cord, although with smaller and slower potentials than the ventrally elicited ones. Several fundamental features of the pathophysiology of a severe acute SCI were reproduced in our experiments, such as: 1) an extensive neuronal loss at the site of injury, with a transient drop in tissue oxygen content; 2) a complete interruption of longitudinal input at the level of impact with a disconnection of the sublesional lumbar cord from both, descending motor commands and ascending afferent input; and 3) a momentary suppression of evoked spinal reflexes. Additionally, our setting highlighted the disappearance of the spontaneous motor activity that is characteristically recorded from the spinal VRs of preparations isolated from newborn rats, showing a drop in the excitability of sensorimotor networks, resembling the flaccid muscle tone displayed in clinics during a spinal shock.

As opposed to the massive neuronal loss at the site of injury, sublesional lumbar motor pools did not undergo any histological changes, while the functionality of locomotor networks was slightly affected, even after two hours from the impact, showing less coordinated locomotor-like cycles, especially from flexor motor pools. Interestingly, after SCI, tail flexor motoneurons underwent distinct morphological alterations, such as a reduction in soma size and an overall decrease in dendritic branching, which concur to the development of spasticity in chronically injured animals (Kitzman, 2005). These morphological changes parallel the selective expression of GABA receptor subunits for flexor, but no extensor, motoneurons in chronic paraplegic rats that with a spinal transection during the first postnatal week (Khristy et al., 2009). Similarly, in paraplegic patients, the appearance of spasms is associated with the exaggerated appearance of flexor reflexes (Hiersemenzel et al., 2000).

In our experiments, the large depolarization triggered by the trauma was further broadened by the low extracellular concentrations of calcium ions, which facilitate the extrusion of calcium from injured spinal cells. This observation confirms the well-known massive calcium release during acute SCIs (Young and Koreh, 1986). Surprisingly, we discovered a crucial role for the rapid outflow of chloride ions in sustaining injury potentials immediately after trauma, as we noticed a maximal depolarization upon increasing the driving force for chloride ions. To the best of our knowledge, this is the first time that chloride ions have been linked to the initial response triggered by a physical trauma to the spinal cord. This evidence could be pivotal in deciphering the origin of the dysregulation of intracellular chloride concentrations that sustain spasticity in persons with chronic SCI.

### A novel in vitro model to trace the immediate consequences of a physical injury to the spinal cord

Starting from the pioneering device introduced by Allen (Allen, 1911), several variations of the original weight drop impactor have been proposed (Wrathall et al., 1985; Kwo et al., 1989) and adopted worldwide, with a standardization for adult rodents (Young and Bracken, 1992; Basso et al., 1996). These resources had a tremendous impact on current knowledge about SCI, but several features of the technique inherently move the outcomes of these experimental injuries far from a clinical scenario. First of all, ethics requires animals to be fully anesthetized, inevitably affecting the composition of spinal tissue (Salzman et al., 1993; Robba et al., 2017; Davis and Grau, 2023). Secondly, breathing chest movements and heartbeat contribute to uniquely vary inter-animal conditions at the time of injury, hence limiting the consistency of the outcome produced by any two identical injury paradigms. Furthermore, the technique only allows a dorsal injury, avoiding the challenging surgical procedures required to transiently move the trunk internal organs to have access to the ventral cord. Hence, it is impractical to replicate a ventral SCI, which is a widely spread condition in clinical epidemiology (Ahuja et al., 2017). Last but not least, currently available impactors generate mechanical interference, coming from the engine or the piston that drive the rod’s vertical displacement, jeopardizing any simultaneous electrophysiological recordings close to lesion site. Our system overcomes all mentioned weaknesses, albeit limited to basic research investigations using in vitro tissue isolated from newborn rodents. Our approach is not intended to replace preclinical tests for the translation of novel treatments in clinics, but to offer an optimal complementary step to challenge innovative basic ideas to target the immediate consequences of a physical injury to the nervous tissue. Our platform is unique in tracing the events at the base of a spinal shock, a topic that has been quite forsaken among the vast SCI research. The constraint of using immature tissue, because of its optimal in vitro survival, hinders highly detailed studies on the physiopathology of adult SCIs, but opens up to the investigation of the still underexplored field of pediatric injuries (Carreon et al., 2004).

In addition, some in vitro models have been defined to explore the mechanical stimulation of the CNS, studying axonal mechanobiology and neuronal membrane deformations, using stretch forces (Aomura et al., 2016) or shear strains (LaPlaca et al., 2005; Bottlang et al., 2007) on cell cultures, and on organotypical or acute CNS slices. While these techniques trace the molecular dynamics at single cell level after mechanical forces have been applied, they cannot pair the informative content of simultaneous electrophysiological recordings. On the other hand, our calibrated microimpactor applies even sudden and orthogonal compressive forces to the nervous tissue of an isolated preparation of the entire CNS, that maintains the multifaced composition of the distinct neural structures. Notably, the low noise design of the device and the stability of the preparation at the impact site allows continuous and stable DC recordings even at the time of impact, with no artifact that prevents signal acquisition after the impact. To the best of our knowledge, no other electrophysiological setups are currently available to record spinal potentials, apart from one attempt that provided recordings after more than four minutes from the impact and after electrodes had been replaced and repositioned, hence limiting the reliability of internal pre-injury controls (Goodman et al., 1985).

### Neuronal source of injury potentials acquired in the current study

The main concern of any well-educated electrophysiologists is the certainty of the genuine biological origin of any acquired signals, to exclude any confounding baseline drifts due to environmental interferences or electromechanical artifacts from the equipment of electrophysiological set-ups. That said, albeit the presence of injury potentials triggered by a physical impact to the cord has already been reported (Goodman et al., 1985; Wang et al., 2015), we wanted to verify the nature of the large depolarization we recorded about 200 ms after the impactor’s functioning. Noteworthy, the device was carefully designed, fabricated, and tested to minimize any sources of electromagnetic emissions, which are stereotyped and instantaneous at the moment of activation.

We collected several convincing proofs about the spinal origin of the potentials we acquired after impact delivery. Namely, we observed that: 1) potentials occur with a latency of hundreds of milliseconds from both engine activation and actual physical strike to the cord; 2) while artifacts are synchronous across all recording sites, the potentials we derived from VRs own different latencies and slower potentials the farther we moved from the impact site; 3) similarly, derived potentials propagate ventro-dorsally, appearing on DRs only after homologous VRs; 4) motor reflexes are suppressed at the top of each potential, and they gradually recover during baseline repolarization, similarly to the reappearance of motor reflexes washing out from high K^+^ concentrations; 5) the injury potential pairs with a reduction in tissue oxygen; 6) amplitude and profile of potentials are modulated in the presence of modified ion solutions.

Additionally, we designed several experimental protocols to confirm that the large deflection of DC level following the impact corresponds to a real neuronal potential, and it is not the mere result of either the engine interference, mechanical movements produced, or the quick displacement of the tip in the bath. Thus, tests aimed at proving that: I) when the device was activated in the bath close to the preparation, but without touching the cord, no baseline drifts were produced; II) when serial impacts of equal severity were applied to the same site, the peaks of potentials remained stable, hence excluding any summation of artifacts; III) no potentials were recorded when the device acted on a preparation that was already maximally depolarized by high K^+^ concentrations; IV) no DC deflections were recorded when the impact was inflicted to a heat-inactivated anoxic spinal cord; V) the injury-related potential was lost when a second impact was inflicted after complete disconnection of the lesion site from the recording VRs. This convincing evidence proves that potentials recorded in correspondence to the activation of the impactor are not artifacts driven by the device engine, nor by the movement of the tip in the recording chamber filled with Krebs medium. Interestingly, the novel platform we introduced allows to elicit and quantify true injury potentials, making it a reliable and consistent tool to study spinal mechanobiology in vitro.

### Chloride surge after a traumatic injury to the cord: a potential link to clinical spasticity

The insurgence of spasticity-like behaviors in SCI rodents has been convincingly attributed to a dysregulation of intracellular chloride concentrations (Boulenguez et al., 2010; Mazzone et al., 2021), with a reduced expression of the membrane carrier KCC2, which co-transports potassium and chloride outside the cell (Boulenguez et al., 2010). However, how an SCI affects chloride exchange in spinal neurons is still to be clarified. The current study suggests that a large chloride conductance sustains the early depolarization that follows a physical injury to the spinal cord. Indeed, the peak of potential was higher for impacts occurring in a low-chloride modified medium, which increases the driving force of inward chloride currents (Takahashi, 1990). We hypothesize that the immediate overflow of chloride ions triggered by a physical injury to spinal tissue sustains a surge of extracellular chloride concentrations, possibly reversing the equilibrium potential of chloride ions. Starting from this early excitatory phase, the net movement of chloride ions across the membrane of spinal neurons possibly still remains perturbed throughout the chronic phase, leading the development of spasticity. Moreover, our results show both, a stable suppression of spontaneous motor activity from VRs and a transient phase of areflexia, right after the impact, followed by a gradual recovery of reflex responses during the following repolarizing phase. However, since motor reflexes consistently reappeared after a fixed amount of time from the impact, regardless of the extent of injury-induced depolarization and of the low concentration of extracellular chloride ions, it can be assumed that reflex depression was not a mere consequence of the overflow of chloride ions triggered by the impact. Thus, network excitation due to the immediate depolarization might be contrasted in the early phases by a following large depression of network excitability that is not directly linked to the extent of the first depolarization. This network inhibition would deserve further pharmacological investigations. Later on, network hyperexcitability prevails and thus spasticity appears. However, time constraints related to our acute in vitro model hindered the possibility to test the occurrence of any chronic spastic-like activity after injury. We are also aware that the neonatal spinal cord used in the current study still presents an immature and opposite reversed chloride gradient (Gao and Ziskind-Conhaim, 1995). This latter feature, although far from adult physiology, makes the neonatal spinal tissue much closer to the extracellular environment after SCI, where the chloride equilibrium is reverted towards that of immature tissues (Lu et al., 2008). We speculate that, albeit spasticity clinically appears only later on, after the recovery from the spinal shock, the molecular elements at the base of spasticity, such as the dysregulation of intracellular chloride concentrations, already start hundreds of milliseconds after the physical injury to the cord. This hypothesis supports the rationale for introducing immediate pharmacological (Liabeuf et al., 2017; Marcantoni et al., 2020) or electrical (Mekhael et al., 2019; Malloy and Côté, 2024) interventions to neuromodulate the shift in chloride concentrations as an early treatment to alleviate the appearance of spasticity in chronic SCIs.

### Transient changes affecting the entire CNS

In the majority of our experiments, we observed that respiratory activity driven by the respiratory central pattern generator (CPG) in the brainstem was temporarily disrupted by spinal cord injury (SCI), presenting a transient pause that suggests an acute supraspinal alteration in network activity. While local effects of SCI have been thoroughly explored across various models, the broader impact on suprapontine structures, particularly within the cerebral cortex, remains less understood. Previous research has demonstrated that SCI can lead to both immediate and long-term reorganization of the cerebral cortex. For instance, an immediate functional reorganization of the primary somatosensory cortex was observed in anesthetized rats following a complete thoracic spinal cord transection (Aguilar et al., 2010). Additionally, SCI has been linked to inflammation in the brain, notably within the primary motor cortex, marked by the activation of microglia (Wu et al., 2014; Hu et al., 2022). Astrocytes can also be relevant to the effects elicited by SCI, as they play a crucial role in shaping neural circuits, including the regulation of synaptic development and function, as well as in neuroinflammatory contexts where they may exhibit both neuroprotective and neurotoxic actions. However, to the best of our knowledge, the immediate effects of SCI on cortical astroglia have not been investigated. Our research aimed to assess the immediate impact of SCI on cortical astrocytes in the primary motor cortex and the primary and secondary somatosensory areas. We observed a significant decrease in astrocyte density in the primary motor cortex during the acute phase, 25 minutes post-impact, with no significant changes detected in the late phase (two hours post-injury). Conversely, a significant decrease in astrocyte density was noted in the primary and secondary somatosensory areas during the late phase, but not earlier. This shift from an early effect in one area to a later effect in another mirrors the changes observed in the spinal cord. Given that our experiments were conducted during the peak period of astrocyte generation in the rodent cortex (P0-P3), the observed changes in astrocyte density may reflect variations in the rate of astrocyte generation. We propose that the decrease in astrocyte density is likely attributable to a slowdown in cortical astrogenesis in response to the spinal injury, rather than a selective loss of astrocytes. This suggests a broader impact of SCI on cortical function and structure beyond the immediate site of injury.

Our findings suggest a complex relationship between SCI and cortical astrocyte responses, with significant changes in astrocyte density occurring in distinct temporal and spatial patterns across different cortical regions. These observations highlight the potential for SCI to influence cortical function and development far beyond the immediate site of injury. The decrease in astrocyte density may signify a disruption in the normal trajectory of cortical astrogenesis in response to SCI during development, potentially impacting cortical plasticity and recovery. Future studies will be necessary to elucidate the molecular mechanisms underlying these changes in astrocytes following SCI, also of different severity, in both early postnatal models and in adult models. Investigations into the specific signaling pathways involved in astrocyte responses could provide valuable insights for targeted interventions to assess astrocyte activity after SCI and hopefully to better diagnose functional recoveries. Additionally, exploring the interactions between astrocytes, other glial cells, and neurons in the context of SCI could further our understanding on the precise roles played by astrocytes in the cortical response to SCI.

## Acknowledgments

GT is grateful to Mrs. Elisa Ius for her excellent assistance in preparing the manuscript and to John Fischetti for technical support in fabricating the impactor. The study was supported by intramural SISSA grants through the 5xMILLE2020 framework.

CF is grateful for Early Career Fellowship awarded to CF from the Human Technopole (Milan, Italy).

## Abbreviations

ANOVA: analysis of variance
C: cervical
CCF: cross-correlation function
Ca^2+^: calcium ions
Cl^−^: chloride ions
CNS: central nervous system
Ctrl: control
CV: coefficient of variation
DAPI: 4’, 6-diamidino-2-phenylindole
DR: dorsal root
DRG: dorsal root ganglia
K^+^: potassium ions
l: left
L: lumbar
P: postnatal
PO2: partial pressure of oxygen
r: right
RMS: root mean square
SCI: spinal cord injury
SD: spreading depression
Th: threshold
T: thoracic
VR: ventral root.

## Author Contributions

GT contributed to the study conception and design. AM, GM, LM, GP and GT performed experiments. Material preparation, data collection and analysis were performed by all authors. The first draft of the manuscript was written and illustrated by GT and AM. GM and CF commented on previous versions of the manuscript. All authors approved the final manuscript.

## Data Availability

The datasets generated during and/or analyzed during the current study are available from the corresponding author on reasonable request.

## Ethics approval

The study was performed in line with the principles of the Italian Animal Welfare Act 24/3/2014 n. 26 implementing the European Union directive on animal experimentation (2010/63/ EU). The study complied with the ARRIVE guidelines.

## Consent to participate

All authors give their formal consent to participate to the present manuscript.

## Consent for publication

All authors give their formal consent for the publication of the present manuscript.

## Supplementary Information

**Supplementary figure 1.**
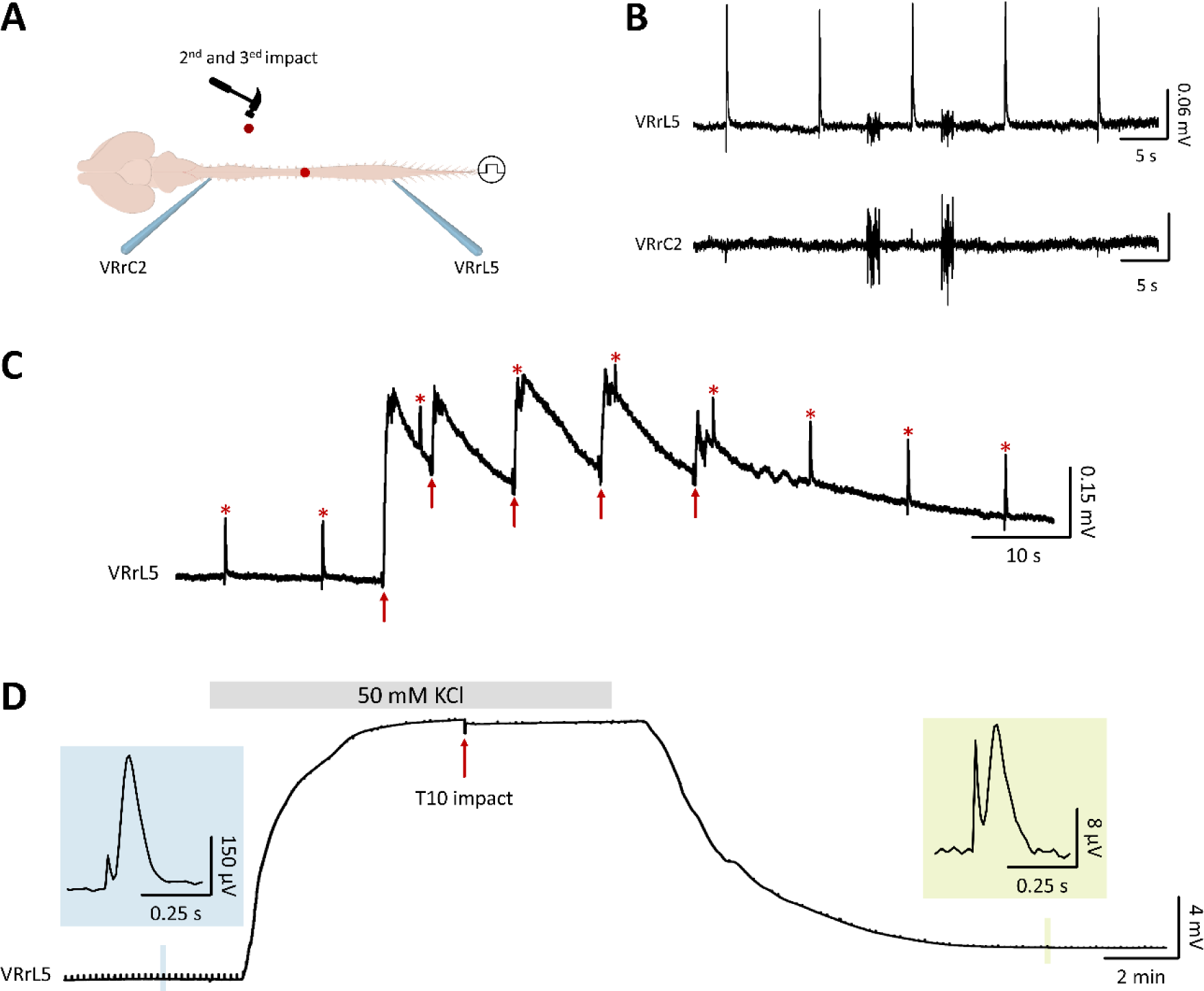
**A.** The cartoon describes an experiment in which, after an initial impact at T10, the tip of the impactor was lowered twice in the recording chamber far from the preparation. Simultaneous VR recordings were taken during the continuous delivery of electric pulses to sacrocaudal afferents. **B.** Two impacts delivered in the bath, away from the preparation, did not depolarize traces from VRrL5 and VRrC2, but generated brief noisier baselines. **C.** A VRrL5 recording during continuous electrical stimulation (red stars) applied to sacrocaudal afferents (0.1 Hz). Five serial impacts were applied at T10 (red arrows) showing consistent depolarization peaks. **D.** An impact (red arrows) delivered at the peak of a large depolarization elicited by 50 mM KCl did not generate any injury-potential depolarizations. After 15 min from the impact in 50 mM KCl, an electrically induced reflex (pale green) reappeared, albeit 95% lower than the peak of a pre-injury control response (pale blue).

**Supplementary figure 2.**
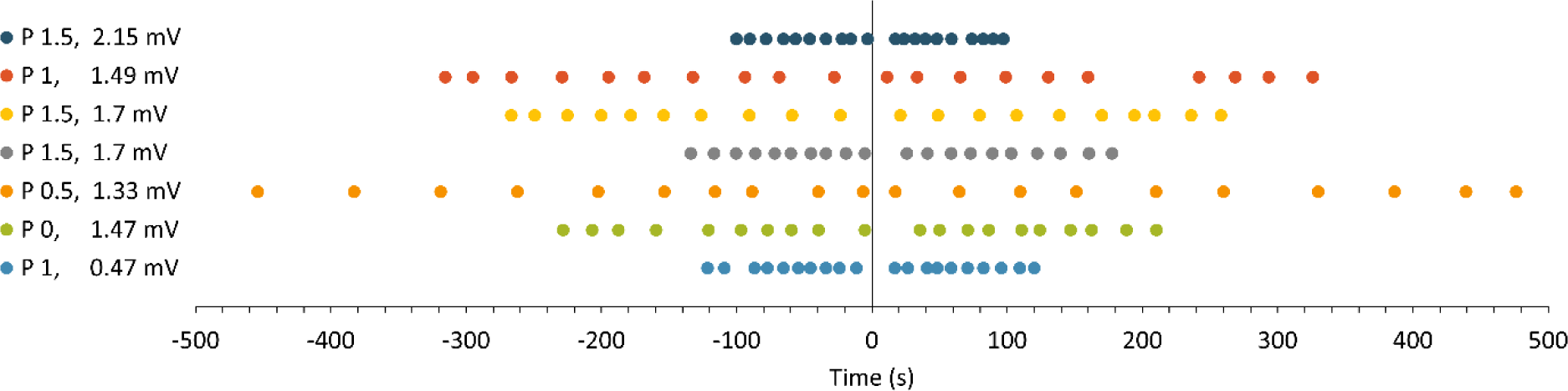
Data pooled from 7 experiments display 20 respiratory events (colored dots) from VRrC2 around the impact (time = 0 s). After the impact, the first respiratory bursts were transiently paused in 4 experiments (blue, green, grey, dark blue). Age of preparations and injury potentials recorded from VRrC2 are both listed on the left side for each experiment.

**Supplementary figure 3.**
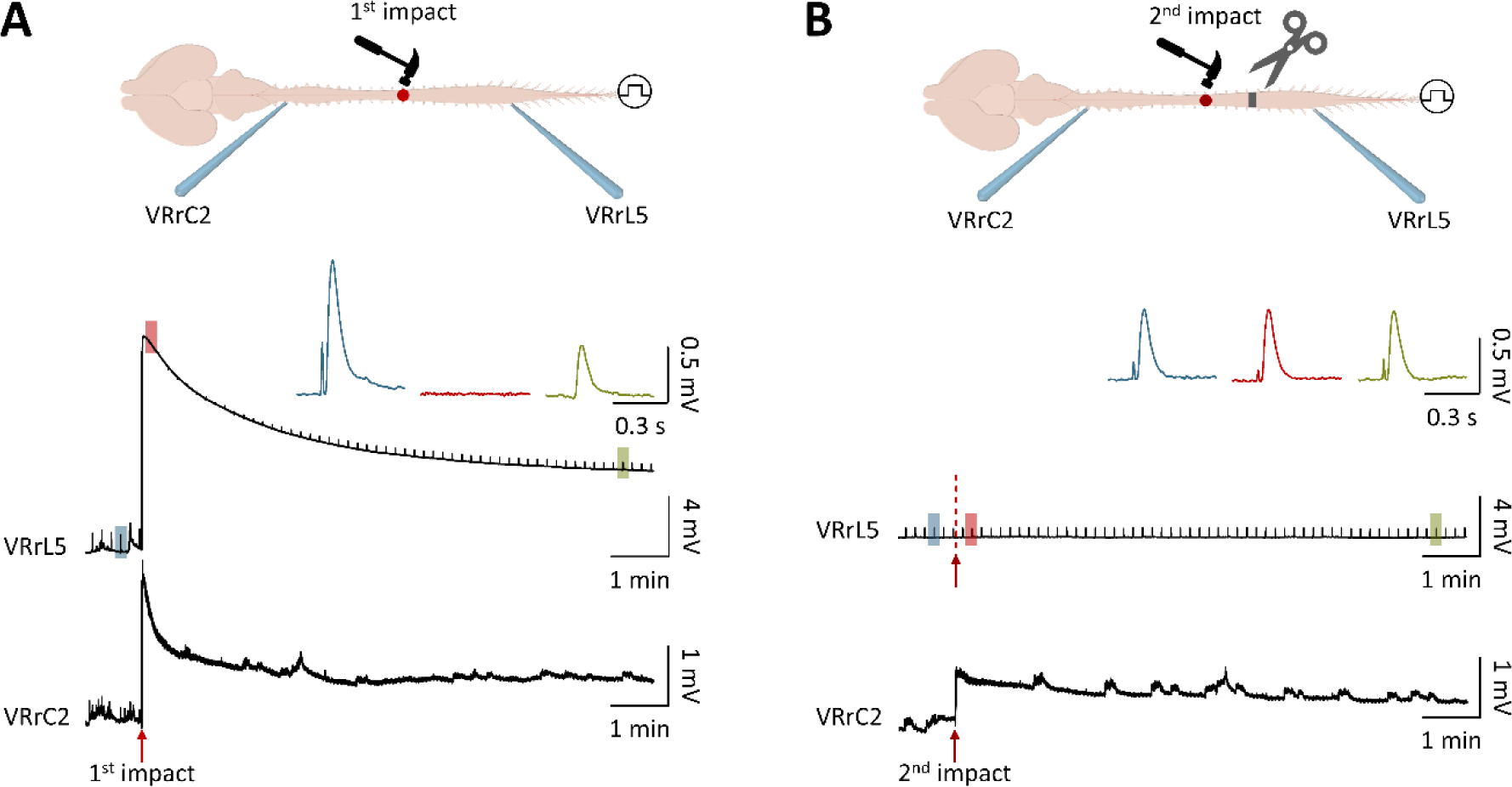
**A.** In the cartoon, glass electrodes derive signals from VRrL5 and VRrC2 from a CNS preparation that was injured at thoracic level (red dot and hammer). Below, simultaneous traces during continuous delivery of electrical pulses to sacrocaudal afferents (0.1 Hz, intensity = 160 µA) around the impact (red arrow) show synchronous injury potentials from VRrL5 and VRrC2 with suppressed reflexes that were reversible for VRrL5 but not for VRrC2. Examples of magnified single reflex responses from VRrL5 are reported as top inserts: in control (blue trace), immediately after the impact (red trace) and after 8.2 min recovery (green trace). Normal reflexes are abolished immediately after the impact, but reappear later on. **B.** In the same preparation, a second impact to the same site, but after a complete transection at L1 spinal level (top cartoon) elicits an injury potential only from VRrC2, while reflexes from VRrL5 were unchanged by the impact to the disconnected rostral cord. Examples of magnified single reflex responses from VRrL5 show the same peak amplitude in control (blue trace), immediately after (red trace) and 8.2 min after (green trace) the impact to the disconnected rostral cord.

**Supplementary figure 4.**
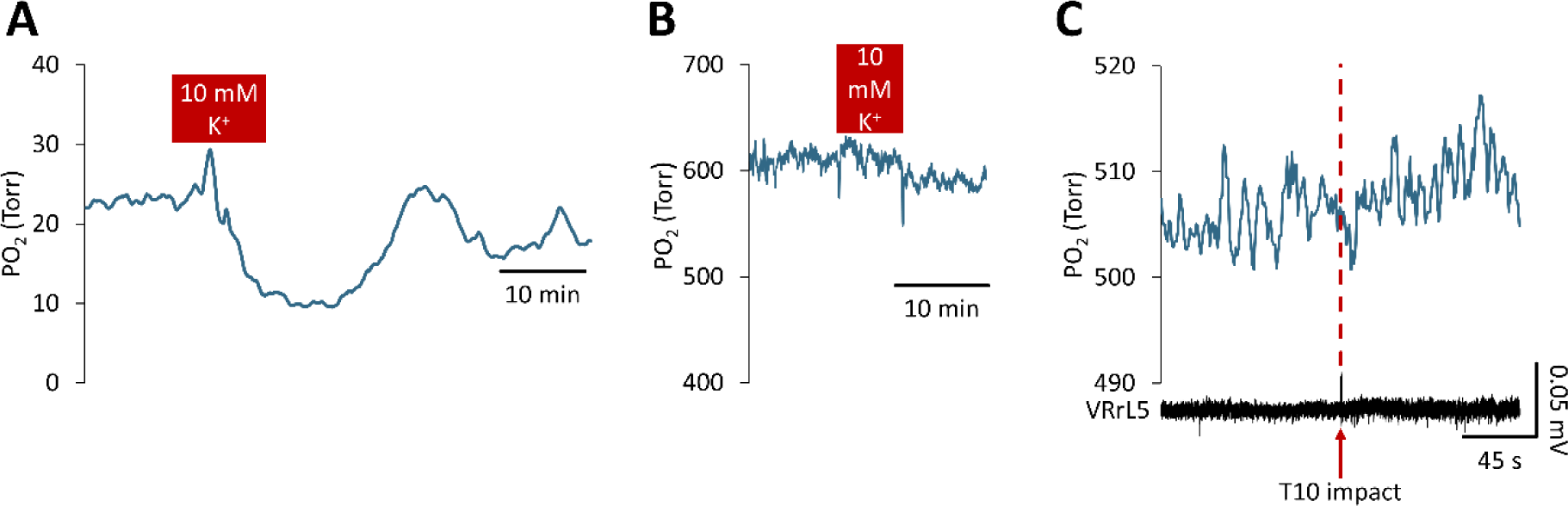
**A.** Time course of average spinal partial pressure of oxygen (PO2) derived at L1 spinal level (n = 8). 10 min of high K^+^ (10 mM, red rectangle) transiently drops the content of tissue oxygen that later recovered. **B.** PO2 assessment in a chamber without any preparation shows perfusion of high K^+^ (10 mM, red rectangle) does not affect the microsensor probe. **C.** Simultaneous tissue oxygen values from L1 spinal level (top) and extracellular recordings from VRL5 (bottom) are taken around the impact at T10 (red dotted line) in a CNS preparation exposed to 100ᵒ C. Unchanged tissue oxygen levels and absence of injury potentials after the impact proves that the CNS preparation is not vital after heat shock.

**Supplementary figure 5.**
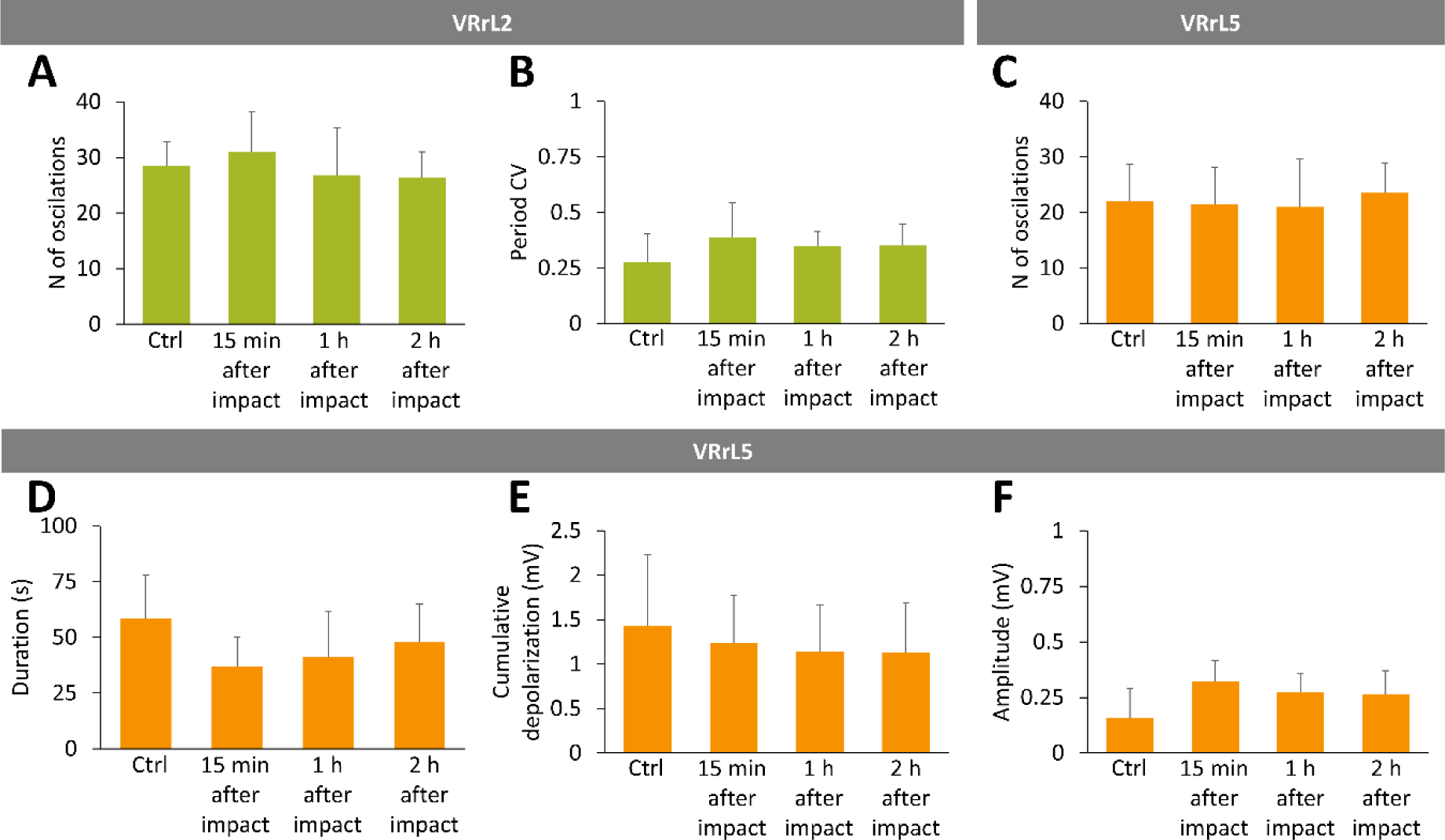
**A-B.** Green bars describe average values for the main descriptors of fictive locomotor patterns recorded from VRrL2 in control and at 15 minutes, 1 hour, and 2 hours following injury. **A.** No changes occur in the number of oscillations. **B.** Coefficient of variation (CV) of period did not change after the impact. **C-F.** Orange bars describe average values for the main descriptors of fictive locomotor patterns recorded from VRrL5 in control and at 15 minutes, 1 hour, and 2 hours following the injury. **C.** Number of oscillations did not vary after the impact. **D.** Duration of oscillations was unchanged post-injury. **E.** No changes occur in the values of cumulative depolarization. **F.** Amplitude of oscillations remained unvaried.

**Supplementary figure 6.**
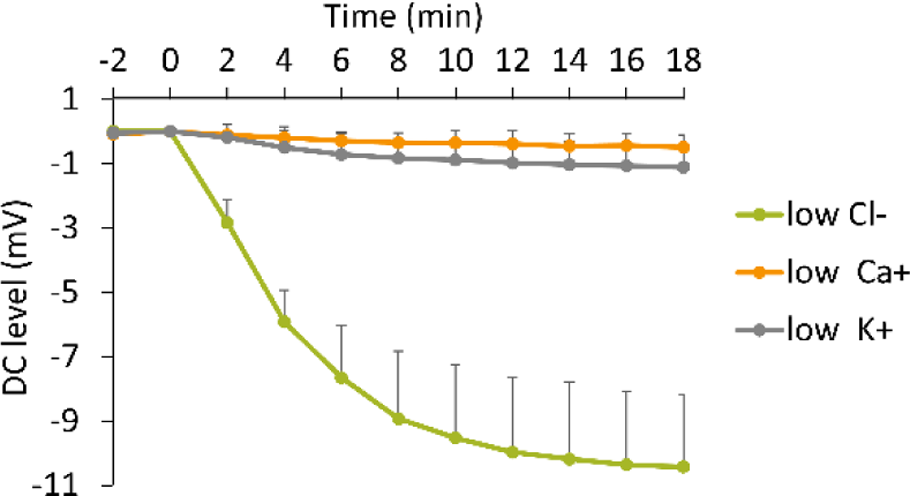
The time course of the DC-level of the VRrL5 baseline during perfusion with modified Krebs solutions for low Cl^−^ (green), low Ca^2+^ (orange) and low K^+^ (grey).

